# inDrops-2: a flexible, versatile and cost-efficient droplet microfluidics approach for high-throughput scRNA-seq of fresh and preserved clinical samples

**DOI:** 10.1101/2023.09.26.559493

**Authors:** Simonas Juzenas, Vaidotas Kiseliovas, Karolis Goda, Justina Zvirblyte, Alvaro Quintinal-Villalonga, Juozas Nainys, Linas Mazutis

## Abstract

The development of a large variety of single-cell analytical techniques has empowered researchers to explore diverse biological questions at the level of individual cells. Among these, droplet-based single-cell RNA sequencing (scRNA-seq) methods have been particularly prevalent owing to their high-throughput capabilities and reduced reaction volumes. While commercial systems have contributed to the widespread adoption of droplet-based scRNA-seq, the relatively high cost impose limitations for profiling large numbers of samples. Moreover, as the scope and scale of single cell sequencing methods keeps expanding, the possibility to accommodate diverse molecular biology workflows and inexpensively profile multiple biospecimens simultaneously becomes highly relevant. Herein, we present inDrops-2: an open-source scRNA-seq platform designed to profile fresh or preserved clinical samples with a sensitivity matching that of state-of-the-art commercial systems, yet at a few folds lower cost. Using inDrops-2, we conducted a comparative analysis of two prominent scRNA-seq protocols – those based on exponential and linear amplification of cDNA – and provide useful insights about the pros and cons inherited to each approach. We showcase the utility of inDrops-2 by simultaneously profiling 18 human lung carcinoma samples, all in one run, following cell preservation, long-term storage and multiplexing, to obtain a multiregional cellular profile of tumor microenvironment. The scalability, experimental flexibility and cost-efficiency offered by inDrops-2 should make it appealing for various single-cell transcriptomic studies.

## Introduction

Comprehensive molecular characterization of biological samples increasingly relies on the accessibility of single-cell technologies [1]. Over the last few years a large array of platforms and methods for single-cell analysis have been introduced, thereby opening a new era of single-cell –omics [2]. In this venture, single-cell RNA-sequencing (scRNA-seq) techniques have been particularly impactful and have gained increasing popularity. Rich biological information encoded in gene expression of individual cells that is captured by scRNA-seq methods have played a critical role, for example, in identifying new cells types in human body [3–5], delineating cancer heterogeneity [6–8] and patient response to therapy [9–11], and have advanced our understanding of various biological systems [12–16]. To this date scRNA-seq represents a leading technology for building cell atlases of the human body and diseases [17–20] and is likely to remain such in a foreseeable future.

Arguably, among scRNA-seq platforms developed to-date [21–28], the most widely used are plate-based and droplet-based systems, each with unique strengths and weaknesses. The plate-based techniques provide an advantage for targeted applications, for instance, when cells of interest are isolated by FACS into microtiter plates for subsequent full-length scRNA-seq, or copy number variation (CNV) analysis [29–31]. These platforms often provide superior sensitivity, although at higher cost and limited throughput. In contrast, droplet-based methods rely on 3’ or 5’ end RNA sequencing and offer a few orders of magnitude higher throughput as well as significantly lower cost. Historically, two droplet-based techniques originally reported side-by-side in 2015 [21, 22] have paved the way for high-throughput single-cell transcriptomics. Built on these innovations commercial systems such as 10x Chromium^TM^ [25] (analogue to inDrops) and Nadia^TM^ [32] (analogue to drop-seq) have followed, providing broad accessibility to scRNA-seq technology.

Commercial systems ensure operational reproducibility and quality, making it a primary choice for single-cell transcriptomics studies. However, profiling single cells at >10^5^ scale using commercially available droplet-based systems, while feasible, can lead to unsustainable financial burden; especially among research groups with limited resources. Open-source systems such as inDrops [22] and drop-seq [21], or their modifications [33–35], offer lower operation costs and can accommodate diverse needs of researchers such as processing unconventional samples [12, 36]. However, open-source systems often exhibit reduced sensitivity (e.g. transcript capture) when compared to commercial peers [37]. Furthermore, barcoding 10^5^-10^6^ individual cells requires extended encapsulation during which the cells of interest may undergo undesirable transcriptional changes. Therefore, in addition to high sensitivity, for scRNA-seq method to be broadly applicable it also needs to ensure the preservation of the native state of cell transcriptome during the course of the workflow.

In this work we present inDrops-2, an open-source droplet microfluidics platform for performing high-throughput scRNA-seq studies of live or fixed cells with the transcript and gene detection similar to that of state-of-the-art commercial platform (10x Chromium v3), yet at the 6-fold lower cost and a throughput of 5,000 cells min^-1^. Using inDrops-2, we implemented two most frequently used scRNA-seq protocols; one based on linear amplification of copy DNA (cDNA) by *in vitro* transcription (IVT), and another based on exponential cDNA amplification by PCR following template switching (TS) reaction. Side-by-side comparison revealed that both scRNA-seq protocols are well-suited for profiling heterogeneous cell populations, yet they also carry important technical differences. The libraries constructed following linear amplification exhibited higher complexity, recovered a higher number of genes, yet the protocol is labor intensive and takes 2 days to complete. The libraries constructed with TS-based approach are simpler to implement and display lower technical variability, yet the sequencing results revealed biased towards capturing shorter genes.

To further expand the applicability of inDrops-2, we report a cell preservation protocol for processing clinical samples comprising as little as 20,000 cells. We show that dissociated cells acquired from clinical specimens can be stored in a dehydrated state for extended periods of time and later multiplexed by covalently conjugating to DNA oligonucleotides [39] for subsequent transcriptomic analysis. We performed multiplexing of methanol-preserved lung carcinoma samples and applied inDrops-2 to obtain a multiregional cellular profile of tumor microenvironment. We captured not only all major specialized lung epithelial and infiltrating stromal and immune cell phenotypes but also patient specific cell populations displaying clinically relevant phenotypes. In summary, we present inDrops-2, a sensitive and cost-efficient scRNA-seq method for capturing clinically relevant cell phenotypes from human specimens that underwent preservation, long-term storage and multiplexing.

## Results

### 1. Optimized inDrops for improved transcript and gene detection

We started our study by reexamining the molecular workflow used in the original inDrops technique [22], with a goal to identify critical parameters that may improve transcript capture and detection. Using a microfluidics setup reported in the past [38] and further detailed in Supplementary Fig. 1 we encapsulated lymphoblast cells (K-562) in 1 nanoliter droplets at occupancy of ∼0.3 together with barcoding hydrogel beads carrying photo-releasable RT primers comprising T7 RNA polymerase promoter (T7p), cell barcode, unique molecular identifier (UMI) and poly(dT19) sequence (Supplementary Table S1). We adjusted the flow rates of the microfluidics platform to achieve the high-throughput of 1 million droplets per hour, while maintaining high hydrogel bead loading at >85%, and stable droplet formation (Supplementary Fig. 1d). Under this set-up, approximately 300,000 single cells can be encapsulated in less than an hour, with a low (∼3%) doublet rate. We prepared scRNA-seq libraries following a workflow outlined in Fig. 1a, while systematically examining each step in the protocol. Specifically, we profiled 1000-3000 cells per condition, by collecting approximately 10,000 droplets, photo-releasing RT primers from the hydrogel beads and performing reverse transcription to produce barcoded-cDNA, followed by second strand synthesis and linear amplification by *in vitro* transcription (IVT). The IVT libraries were fragmented with Lewis acid (Zn^2+^ ions), transcribed into single-stranded cDNA and PCR-amplified with sequencing adapters to obtain final gene libraries compatible with Illumina sequencers. After a series of optimizations, we arrived at inDrops-2 (IVT) protocol that showed markedly improved transcript and gene detection, and is provided as Supplementary Protocol 1. The most important findings can be summarized as follows. Compared to original inDrops [22], the barcoded-cDNA material purification and primer dimer removal by solid-phase reversible immobilization (SPRI), rather than digestion with nuclease cocktail, had the most significant impact (Fig. 1b). At sequencing depth of 15,000 reads per cell, inDrops-2 showed increased UMI detection by 2.72-fold (mean ± s.d, 8166 ± 350 vs 2999 ± 461 UMIs, t-test, P_FDR_ < 1 L 10^-300^) and gene capture by 1.86-fold (mean ± s.d., 3399 ± 143 vs 1829 ± 207 genes, t-test, P_FDR_ < 1 L 10^-300^). Replacing SuperScript III with Maxima H-minus reverse transcriptase increased UMI and gene capture further by 1.19-fold and 1.18-fold, respectively (t-test, UMI P_FDR_ = 1.62 L 10^-281^, gene P_FDR_ = 5.82L10^-283^). The improved UMI/gene detection using inDrops-2 was mirrored in primary cells (Supplementary Fig. 1e-g) and was not due to preference for a specific RNA biotype (Supplementary Fig. 1h), and overall displayed lower technical variability (Fig. 1c). The inDrops-2 libraries prepared with SuperScript IV (SS-IV) enzyme displayed slightly higher UMI count (SS-IV 9108 ± 198 vs Max 9108 ± 709, t-test, P_FDR_ = 6.22 L 10^-317^) (Fig. 1d), however, given the significantly higher cost of the SS-IV enzyme, and relatively mild improvements, it was excluded from our subsequent efforts. The inDrops-2 conducted at 42 °C tend to show slightly higher UMI/gene counts than corresponding reaction at 50 °C, irrespectively of the RT enzyme tested (Fig. 1d). No significant effect on UMI/gene recovery was observed when using different commercially available second strand synthesis or *in vitro* transcription kits, indicating that the critical steps for obtaining high UMI counts are indeed mainly related to reverse transcription reaction and subsequent purification of barcoded-cDNA molecules. In summary, single cell transcriptional profiling with inDrops-2 shows markedly improved UMI/gene detection, nearly 20-fold higher throughput (275 cells s^-1^ vs 15 cells s^-1^), and higher quality data (Fig. 1e-g). The step-by-step protocol incorporating aforementioned improvements is accompanying this manuscript as Supplementary Protocol 1.

**Figure 1.**
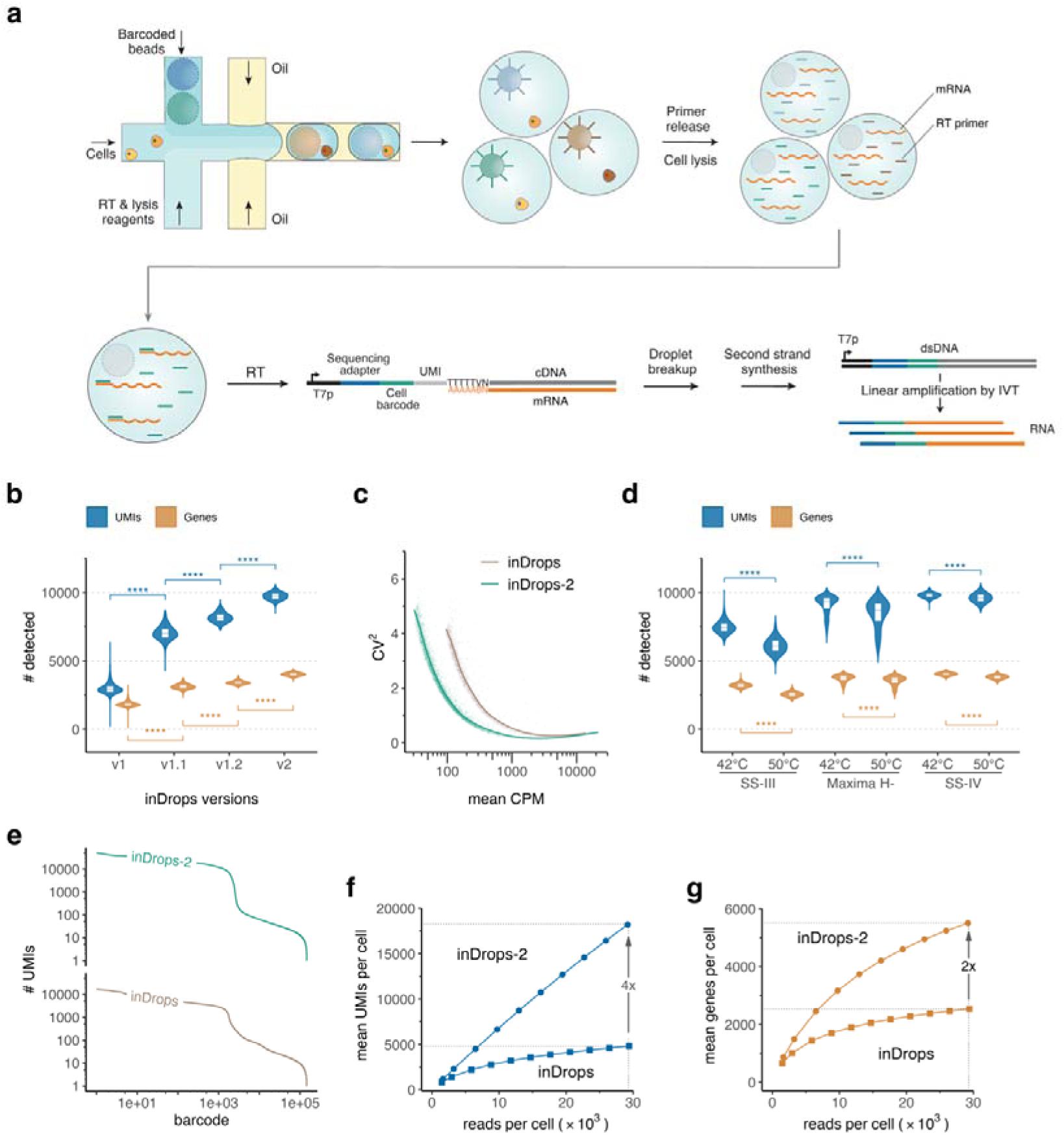
The overview of inDrops-2 performance. **a**) Schematics of inDrops-2 technique using linear amplification by *in vitro* transcription (IVT) of barcoded cDNA. **b)** Detection of transcripts (unique molecular identifiers, UMIs) and genes in K-562 cell line, using different versions of inDrops (see Methods). To normalize for the sequencing depth, each cell barcode was downsampled to 15’000 raw reads. **c)** Comparison of technical variability between inDrops and inDrops-2, where CPM and CV^2^ refers to counts per million and squared coefficient of variation, respectively. **d)** Evaluation of reverse transcription enzymes (SuperScript III Reverse Transcriptase [SS-III], Maxima H minus Reverse Transcriptase [Maxima H-] and SuperScript IV Reverse Transcriptase [SS-IV]) and temperature (42 and 50 _) on the UMI and gene detection. Downsampled to 15’000 raw reads. **e)** Barcode rank plots derived from scRNA-seq data acquired with inDrops-1 and inDrops-2 techniques. Barcodes having at least one UMI are arranged from the highest to the lowest UMI counts. **f)** Mean UMI count per cell as a function of sequencing depth. **g)** Mean gene count per cell as a function of sequencing depth. **b)**, **d)** Boxplots within density violins show median (center line), first and third quartiles (lower/upper hinges), 1.5× interquartile range (lower/upper whiskers); ****P value < 0.0001 (two-sided t-test, Benjamini-Hochberg correction).

### 2. inDrops-2 based on template switching reaction for rapid construction of sequencing libraries

While improved inDrops-2 based on linear amplification enables a substantially higher UMI and gene recovery per single cell, an alternative scRNA-seq strategy commonly used nowadays relies on a template switching (TS) reaction driven by the RT enzyme, and often referred as SMART [39–41]. Owing to the intrinsic terminal transferase activity, the RTase tends to add a few non-templated nucleotides (predominantly cytidines) on the 3’ end of cDNA, and does so preferably if the RNA template is G-capped [42–44]. A large variety of scRNA-seq methods have exploited this unique RT enzyme feature to incorporate the PCR adapters at 5’ mRNA end, and facilitate library preparation for next generation sequencing [25, 29, 40, 45]. Motivated by these and other reports, we sought to adopt a TS-based reaction with inDrops and to compare the UMI/gene capture to those obtained by linear amplification. To differentiate the two scRNA-seq approaches, we refer to them as inDrops-2 (IVT) and inDrops-2 (TS).

At first, we aimed to maximize the barcoded-cDNA yield using inDrops-2 (TS) approach (Fig. 2a). For that purpose, we encapsulated K-562 cells together with barcoding hydrogel beads and RT/lysis reaction mix supplemented with template switching oligonucleotide (TSO), that is required for barcoded-cDNA amplification by PCR (see Materials and Methods). We thoroughly optimized each step of the workflow, namely; cell lysis and RT reaction, TSO concentration, temperature, barcoded-cDNA purification, library fragmentation, A-tailing and adapter ligation, and other parameters (Supplementary Fig. 2) to obtain a robust and reproducible scRNA-seq protocol that is described in details as Supplementary Protocol 2. Next, we evaluated the sensitivity of inDrops-2 (TS) by profiling human PBMCs (ATCC) and compared the results to those obtained with a commercial analogue (10X Genomics, Chromium v3 chemistry). Sequencing results presented in Fig. 2b revealed that at the same sequencing coverage the inDrops-2 (TS) detects nearly the same UMI and gene count in single cells as the current gold-standard in the field. The cell types comprising the biospecimen that were identified with two techniques showed almost identical cell composition (Fig. 2c and 2d) and displayed high correlation in gene expression among respective cell types (Fig. 2e). Furthermore, deep sequencing (mean reads per cell: ∼221,000) unveiled inDrops-2 (TS) to be sensitive enough to detect up to ∼140,000 UMIs (mean: 39,388) and up to ∼7,500 genes (mean: 4,909) in murine lung adenocarcinoma cell line [46] at 0.81 sequencing saturation (Fig. 2f and 2g). Altogether these results reassured us that inDrops-2 (TS) can serve as a highly efficient method for profiling cells of different origin in a cost-effective manner.

**Figure 2.**
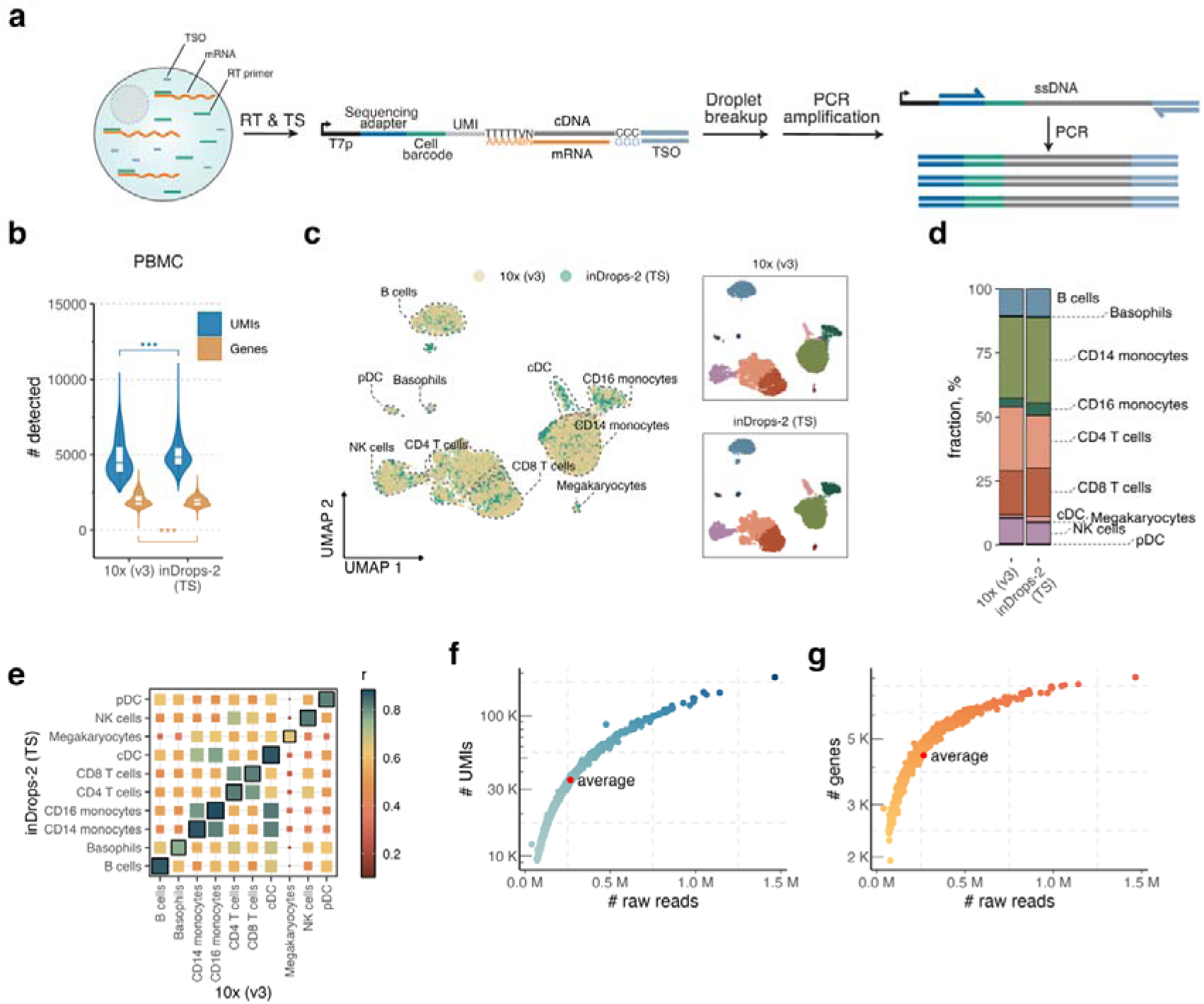
The overview of inDrops-2 using template switching approach. **a**) Schematics of inDrops-2 (TS) technique based on exponential cDNA amplification following template switching (TS) reaction. **b)** Comparison of transcript (unique molecular identifiers, UMIs) and gene detection in human peripheral blood mononuclear cells (PBMCs) between 10X Genomics (v3) and inDrops-2 (TS) platforms at sequencing depth of 20’000 reads per cell. Boxplots within density violins show median (center line), first and third quartiles (lower/upper hinges), 1.5× interquartile range (lower/upper whiskers); ***P value < 0.001 (two-sided t-test, Benjamini-Hochberg correction). **c)** Dimensionality reduction (UMAP) of human PBMCs profiled with 10X Genomics (v3) [n = 4803] and inDrops-2 (TS) [n =6025] and colored by platform (left panel) and annotated cell type (right top and bottom panels). The UMAP representation is based on 3000 highly variable genes after data integration by Harmony (see Methods). **d)** Comparison of cell type fractions recovered with 10X Genomics (v3) and inDrops-2 platforms. **e)** Pearson’s correlation analysis between highly variant genes in cell type clusters (cluster ellipsoids within percentile of 0.5) inferred by inDrops-2 (TS) [rows] and inferred by 10X Genomics (v3) [columns] platforms. **g)** and **h)** Deep sequencing results of scRNA-seq libraries prepared using inDrops-2 (TS) approach. Murine KP cell line was used as a model system. The plots show raw reads of a given cell barcode on the x-axis and UMI **(g)** as well as gene **(h)** counts on the y-axis in a log_10_ scale.

### 3. Comparison of scRNA-Seq protocols based on linear amplification vs. exponential amplification of cDNA

Having established inDrops-2 (TS), we then asked which scRNA-Seq approach, IVT-based or TS-based, can deliver a higher number of unique transcripts and genes. While previous benchmarking studies indicated that scRNA-Seq libraries constructed by linear amplification (e.g, CEL-Seq2) recovers higher diversity of genes as compared to PCR-based approaches such as SMART-seq2 [47], yet the head-to-head comparison of IVT-based or TS-based technique, to the best of our knowledge, is lacking. To perform such a comparison, we first formed an emulsion comprising approx. 5000 cells, compartmentalized along with hydrogel beads carrying barcoding RT primers with T7 RNA polymerase promoter sequence, and 25 µM TSO. Upon completion of cDNA synthesis at 42 °C for 90 min, the emulsion was split into two equal fractions and processed separately following either inDrops-2 (IVT) or inDrops-2 (TS) protocol (Fig. 3a). Following this strategy, we prepared and sequenced lung carcinoma and lymphoblast cells. After down-sampling the sequencing depth of each library to 20’000 raw reads per cell, we find both approaches recovering a similar number of UMIs per cell (Fig. 3b). However, a significant fraction of transcripts in TS-based libraries corresponded to the ribosomal protein (RP) genes (Fig. 3c), which are often excluded from downstream analyses. Removing RP genes enhanced the differences between the two protocols with the libraries constructed by linear amplification revealing ∼25% higher UMI and gene detection (Supplementary Fig. 3a). Interestingly, the inDrops-2 (IVT) was also better at capturing unspliced RNAs as confirmed by higher fraction of reads aligning to introns as well as 3’ UTRs, which play a role in post-transcriptional gene expression regulation (Fig. 3d). As a drawback, the IVT-based approach exhibited slightly higher technical noise as well as higher run-to-run variability (Supplementary Fig. 3b). The TS-based libraries exhibited a slightly higher GC content (Supplementary Fig. 3c), in accordance with a higher fraction of coding sequences [48]. Additionally, as opposed to sharp enrichment at the 3’ end, the gene body coverage in the TS-based approach was shifted, implying a trend towards shorter genes (Fig. 3e). Indeed, a striking difference between two scRNA-Seq protocols became evident, when all detected genes were binned according to their length (Fig. 3f-g). This analysis clearly revealed the bias of template-switching based approach towards shorter genes; a trend that was reproduced in independent experiments on PBMCs as well as using 10X genomics (v3) platform (Supplementary Fig. 3d-g). Some of the observed differences could be attributed to an increased probability of stalling and drop-off events by the reverse transcriptase while processing long RNA templates (Supplementary Fig. 3h), to which the IVT-based approach is less sensitive, since the truncated cDNAs can still be linearly amplified and then sequenced. Also, cDNA products corresponding to the internal priming and the transcripts that may lack a G-cap and A-tail such as non-coding RNAs, were more frequently found in inDrops-2 (IVT) libraries (Supplementary Fig. 3i). Therefore, single cell transcriptome libraries prepared with TS-based, or IVT-based, approach are prone to specific technical biases that, when unaccounted, could negatively impact the interpretations of gene expression dynamics in individual cells, as shown by gene set enrichment analysis performed on differentially expressed genes (Fig. 3h and Supplementary Tables 2-3). On another side, both approaches provide high confidence for identifying different cell types, making them suitable for cellular composition profiling and tissue atlases efforts (Supplementary Fig. 3j).

**Figure 3.**
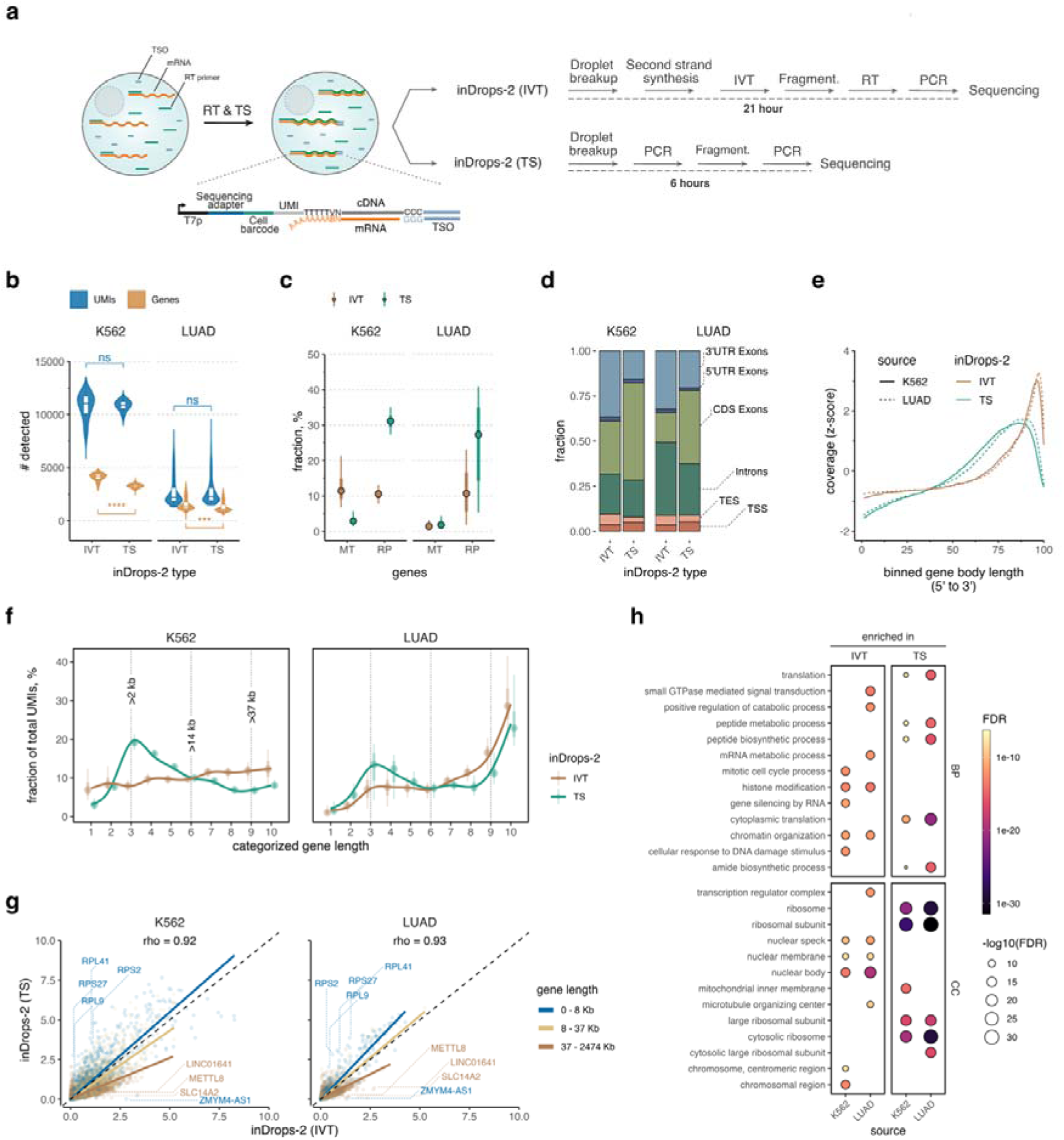
Comparative analysis of scRNA-seq libraries prepared with linear and exponential amplification of cDNA in primary and cultured cells. **a**) Schematics of the experiment. A droplet-based reverse transcription (RT) reaction is performed in the presence of barcoded RT primers (comprising T7 promoter) and template switching oligonucleotide (TSO). The post-RT emulsion droplets were split in two equal fractions and sequencing libraries prepared according to inDrops-2 (IVT) and inDrops-2 (TS) protocol. **b)** Number of UMIs and genes detected in lymphoblast (K-562) and primary lung adenocarcinoma (LUAD) cells in scRNA-seq libraries prepared by linear amplification of cDNA using *in vitro* transcription (IVT) reaction, and by exponential amplification of cDNA following template switching (TS) reaction. Sequencing depth was normalized to 20’000 raw reads per cell. Boxplots display median (center point), first and third quartiles (lower/upper hinges), 1.5× interquartile range (lower/upper whiskers); ****P value < 0.0001, ***P value < 0.001, ^ns^P value > 0.05 **c)** Fraction of genes corresponding to mitochondrial (MT) and ribosomal proteins (RP) in IVT-based and TS-based scRNA-seq libraries. **d)** Fraction of reads mapping to different regions of a gene. **e)** Sequencing coverage across the gene body. To adjust for the sequencing depth, raw coverage values were scaled and centered to a z-score. **f)** Fraction of UMIs as a function of binned gene length. Fractions per cell, alongside with the boxplots, are displayed as curves fitted using loess smoothing and coloured by the inDrops-2 protocol type. **g)** Spearman’s correlation analysis between inDrops-2 (IVT) and inDrops-2 (TS) protocols. Correlation coefficient (rho) is depicted at the top of the scatter plot. Each dot represents size factor-normalized and log2(x+1)-transformed expression levels of detected genes. Dashed line (diagonal) divides panels in two equal parts, whereas blue, yellow and brown lines display linear regression curves corresponding to short (0 – 8 Kb), medium (8 – 37 Kb) and long gene length (37 – 2474 Kb) categories, respectively. Detection of transcripts encoded by longer genes (brown) is improved in IVT-based approach as compared to TS-based scRNA-seq libraries. Note, the median length of protein-coding gene in humans is 26 Kb. Top differentially expressed genes (lowest P value, n=20 in each cell type, based on MAST) that overlap in both primary and cultured cells are annotated using gene symbols. **h)** Gene set enrichment analysis performed on ordered gene list by the level of differential expression (log2 fold change) between IVT-based and TS-based scRNA-seq libraries. Gene ontology (GO) terms BP and CC refers to biological process and cellular compartment, respectively. FDR adjusted P-values are presented as color gradient as well as are shown proportionally to dot size in –log_10_ scale. **f)**, **g)**, Gene categories were determined by binning all genes based on their length into groups with approximately equal number of genes per category.

### 4. Cell preservation for long-term storage and transcriptomic studies of primary cells

Even with high-throughput capabilities offered by droplet microfluidics technology, the barcoding of increasingly high numbers of single cells (>10^5^-10^6^ scale) may require a sufficiently long (>30 min) encapsulation times during which there is an increased risk that live cells will alter their native transcriptional state. To mitigate such risk, it is desirable to safeguard cellular transcriptome by fixing the cells so that no transcriptional changes would occur during the encapsulation process. In this regard, cell preservation in methanol represents an appealing option as it was shown to retain transcriptional signatures of cells, and to be compatible with droplet microfluidics methods [49–51]. Unfortunately, our attempts to adopt aforementioned cell preservation protocols were unsatisfying for PBMCs as we witnessed high variability of UMI / gene capture between the individual runs, and experienced a significant RNA degradation and cell loss due to clumping (Supplementary Fig. 4a). Primary cell recovery was particularly problematic when handling clinical samples comprising a low number (n ≤ 100,000) of cells. We reasoned that excessive centrifugation force that is required to pellet the cells during rehydration might be causing cell clumping and damage. Accordingly, after a series of independent tests we found that methanol-fixed cells placed on 0.65 µm pore size filters (see Supplementary Protocol 3) can be effectively rehydrated without applying an excessive centrifugation (see Methods). We confirmed that RNA integrity of cells in a rehydration buffer containing citrate remains high (RIN > 8) after 30 minutes on ice (Supplementary Fig. 4b) and it is not affected by preservation time (up to 30 days) in methanol (Supplementary Fig. 4c). Importantly, using rehydration columns cell clumping was negligible (Supplementary Fig. 4d), cellular morphology was consistently retained (Supplementary Fig. 4e) thereby increasing the reproducibility of cell recovery.

We then applied inDrops-2 to compare UMI and gene capture of the methanol-preserved vs fresh human PBMCs (Fig. 4). To avoid the RT reaction inhibition by citrate that is present in the rehydration buffer we adjusted the flow rates such that the final dilution of rehydration buffer in a droplet would correspond to 0.1X, or 1.5 mM sodium citrate (see Methods). In addition to benchmarking fresh and methanol-fixed human PBMCs we also tested bone marrow CD34 positive (CD34^+^) cells. Sequencing results presented in Fig. 4a confirmed the efficient transcript recovery in fixed cells by inDrops-2, closely matching those of live PBMCs profiled alongside with commercial platform (10X Chromium V3). As expected, live cells displayed a higher fraction of mitochondrial genes (Fig. 4b), indicative of the undergoing transcriptional response during cell handling procedures. Average gene expression levels exhibited high correlation (Fig. 4c), with gene mapping characteristics matching closely for both fresh and fixed samples (Supplementary Fig. 4f). Performing feature selection, dimensionality reduction and clustering for PBMCs revealed all expected cell populations, including CD4 T, CD8 T, NK, CD14 and CD16 monocytes, dendritic cells and megakaryocytes with similar cell proportions in fixed and fresh samples (Fig. 4d-e). Expression levels and detection rates of PBMC marker genes revealed no noticeable ambient RNA contamination in methanol-fixed cells and resembled those of freshly profiled libraries (Supplementary Fig. 4g). A transcriptional map of CD34^+^ cells reconstructed hematopoietic stem cell differentiation into all known progenitor lineages (Fig. 4f) and, based on marker gene expression, recapitulated the expected trends in hematopoiesis (Fig. 4g). All major lineages, such as common lymphoid progenitors, erythroid precursor cells, megakaryocytes, neutrophil-myeloid progenitors, dendritic cell precursors and mast cell precursors, were present and was cell type composition in agreement between fixed and fresh samples (Fig. 4h). As a final quality measure, we compared mapping statistics and observed no significant differences (Supplementary Fig. 4f). Overall, these results demonstrate that column-based rehydration of methanol-preserved cells ensures: i) minimal cell loss during handling, ii) high number of singlet recovery, iii) efficient UMI/gene detection, iv) accurate recapitulation of the transcriptional signature matching that of live cells, and v) alleviates the adverse effects of cell viability decline (i.e., increased mitochondrial gene expression), caused by extended workflows.

**Figure 4.**
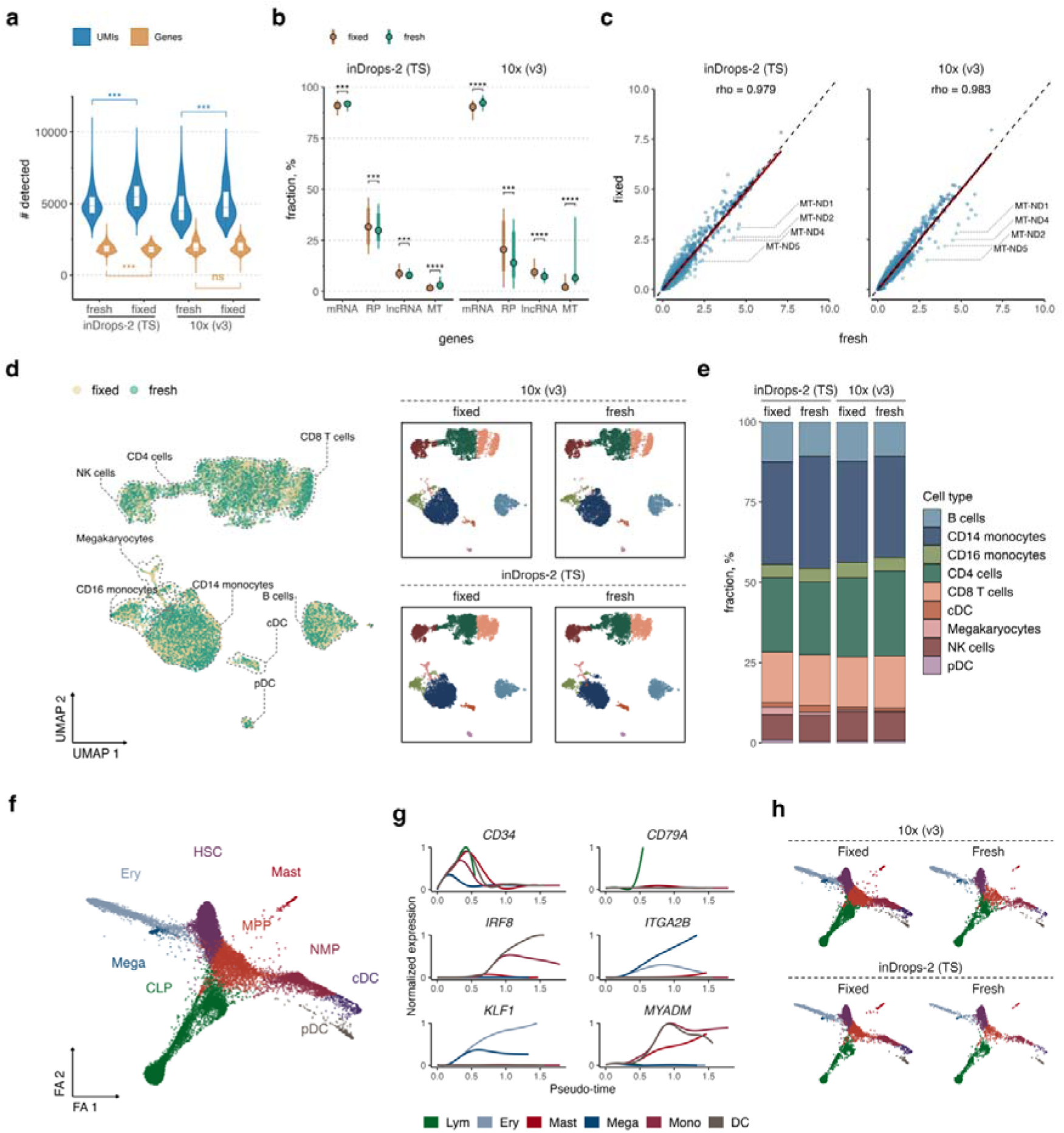
scRNA-seq of fresh and methanol-fixed PBMCs and CD34^+^ cells using inDrops-2 and 10X Genomics (v3) platforms. **a**) UMI and gene detection in fresh and methanol-fixed PBMCs. **b)** Fraction of UMIs along RNA biotypes. **c)** Spearman’s correlation analysis between fresh and methanol-fixed PBMCs sequenced with inDrops-2 and 10X Genomics (v3) platforms. Correlation coefficient (rho) is shown at the top of the scatter plot. Size factor-normalized and log2(x+1)-transformed gene expression levels are displayed as dots. The black dashed line represents diagonal, while the red solid line displays fitted linear regression. **d)** UMAP of fresh and methanol-fixed PBMCs sequenced with inDrops-2 (TS) [n=12279] and 10X Genomics (v3) [n=9582], based on 3000 highly variable genes, and integrated using Harmony. The UMAP is colored by cell preservation type (left panel) and annotated cell type, faceted by platform and preservation type (right top and bottom panels). **e)** Cell types and their fractions recovered in fresh and fixed PBMC samples in inDrops-2 (TS) and 10X Genomics (v3) methods. **f)** Force Atlas (FA) embedding of human CD34^+^ profiled with 10X Genomics (v3) [n=10343] and inDrops-2 (TS) [n=5029] coloured by annotated cell type (HSC – hematopoietic stem cells, MPP – multipotent progenitors, Ery – erythroid progenitors, Mega – megakaryocytes, CLP – common lymphoid progenitors, NMP – neutrophil-myeloid progenitors, cDC – conventional dendritic cells, pDC – plasmacytoid dendritic cell, Mast – Mast cells). The FA is based on 2000 highly variable genes with data integration by Harmony. **g)** Gene expression trends for characteristic lineage genes along Palantir pseudo-time. Genes selected for each lineage were *CD34* for HSC, *CD79A* for CLP, *IRF8* for DC cells, *ITGA2B* for megakaryocytes, *KLF1* for erythroid cells and *MYADM* for myeloid cells. h) FA from f) colored by annotated cell type and faceted by platform as well as cell preservation method. a), b) median (center line or dot), first and third quartiles (lower/upper hinges), and 1.5× interquartile range (lower/upper whiskers); ****P value < 0.0001, ***P value < 0.001, ^ns^P value > 0.05.

### 5. Scaling inDrops-2 with click chemistry hashtags

Sample multiplexing with hashtags provides an appealing option to increase the scale of scRNA-Seq experiments [52–56]. Taking advantage of methanol-based cell preservation, we applied multiplexing by methyltetrazine-modified DNA oligonucleotides or ‘ClickTags’ [52]. The distinct feature of ClickTags is that they do not rely on specific cell epitopes for labeling and can be chemically attached to cellular proteins by Diels–Alder reaction, thus making them applicable to a broad range of cells and biospecimens. We sought to profile human lung tumor microenvironment, by conducting multiregional cell composition and gene expression analysis. The surgical samples acquired from lung carcinoma patients were cut into 3 parts, dissociated and sorted into CD45 positive and CD45 negative compartments [57]. Following FACS, the single cell suspensions were preserved in methanol and transferred to –80 °C for long-term storage. After 4 weeks, the cells were retrieved and while in methanol hashed with ClickTags (see Methods). In total, 18 samples were hashed, then pooled, and following rehydration (see Supplementary Protocol 3) processed according to inDrops-2 (TS) workflow.

After sequencing, filtering and quality control steps (see Methods) we obtained 32,937 high quality cells with a rather consistent number of cells across hashtags (Fig. 5a and Supplementary Fig. 5a). The average UMI and gene count were high, 6959 and 1966, respectively (Fig. 5b and 5c) and matched closely the sequencing statistics of fresh tumor samples (e.g. 5000 UMIs and 2100 genes, on average, at 40,000 reads per cell) [58]. As expected, there was a dependency of ClickTag and UMI counts per cell (Fig. 5d). Following data normalization, feature selection, dimensionality reduction, clustering and visualization with uniform manifold approximation and projection (UMAP) cells were manually annotated using canonical gene markers (Fig. 5e-h). UMAP colored by FACS-sorting label showed clear separation of CD45 positive and negative cells, as expected (Fig. 5g). We detected all major specialized lung epithelial and infiltrating stromal and immune cell phenotypes featuring lung carcinoma disease (Fig. 5e, h), coinciding with previous reports [7, 59, 60]. As it is common to the tumors, there was a high inter-patient variability with regards to the cellular composition (Fig. 5f), while inter-regional differences within individual tumors were not as pronounced (Fig. 5i). High resolution analysis in the non-immune cell compartment uncovered lung-specialized epithelial cells such as alveolar epithelial cells (AEC, markers *SFTPA1, HOPX*), club (*SCGB1A1, SCGB3A2*), ciliated (*CAPS, PIFO*), neuroendocrine (*CALCA, UCHL1*) and basal (*KRT17, KRT15*) cells, as well as patient specific *SPINK1*^high^ club cell population and *MMP7*^high^ alveolar epithelial phenotype (Fig. 5e, j). Recently, it has been reported that *SPINK1* upregulation leads to adverse outcomes in a multitude of cancers, and enhances proliferation and invasion of tumor cells *in vitro* [61]. Interestingly, the *SPINK1*^high^ club cells (*SCGB3A1, SCGB3A2*) in our dataset also expressed distal lung marker *NAPSA* and *CEACAM6* (Supplementary Fig. 5), whereas the latter has been implicated in lung cancer progression and poor clinical outcomes [62]. Another interesting finding was two distinct phenotypes of alveolar epithelial cells – both of them expressing canonical AEC markers (i.e. *SFTPA1, SFTPA2, SFTPC*), yet only one population was marked by *MMP7* and *PRSS2* expression (Fig. 5j). *MMP7* is a widely used biomarker for pulmonary fibrosis, while *PRSS2* is associated with invasive and metastasis promoting features [63]. The co-expression of *PRSS2* and *MMP7* in transitional state epithelial cells has been implicated in idiopathic pulmonary fibrosis [64], thus, our results suggest that *MMP7*^high^ alveolar epithelial cell phenotype might be involved in disease progression and plasticity.

**Figure 5.**
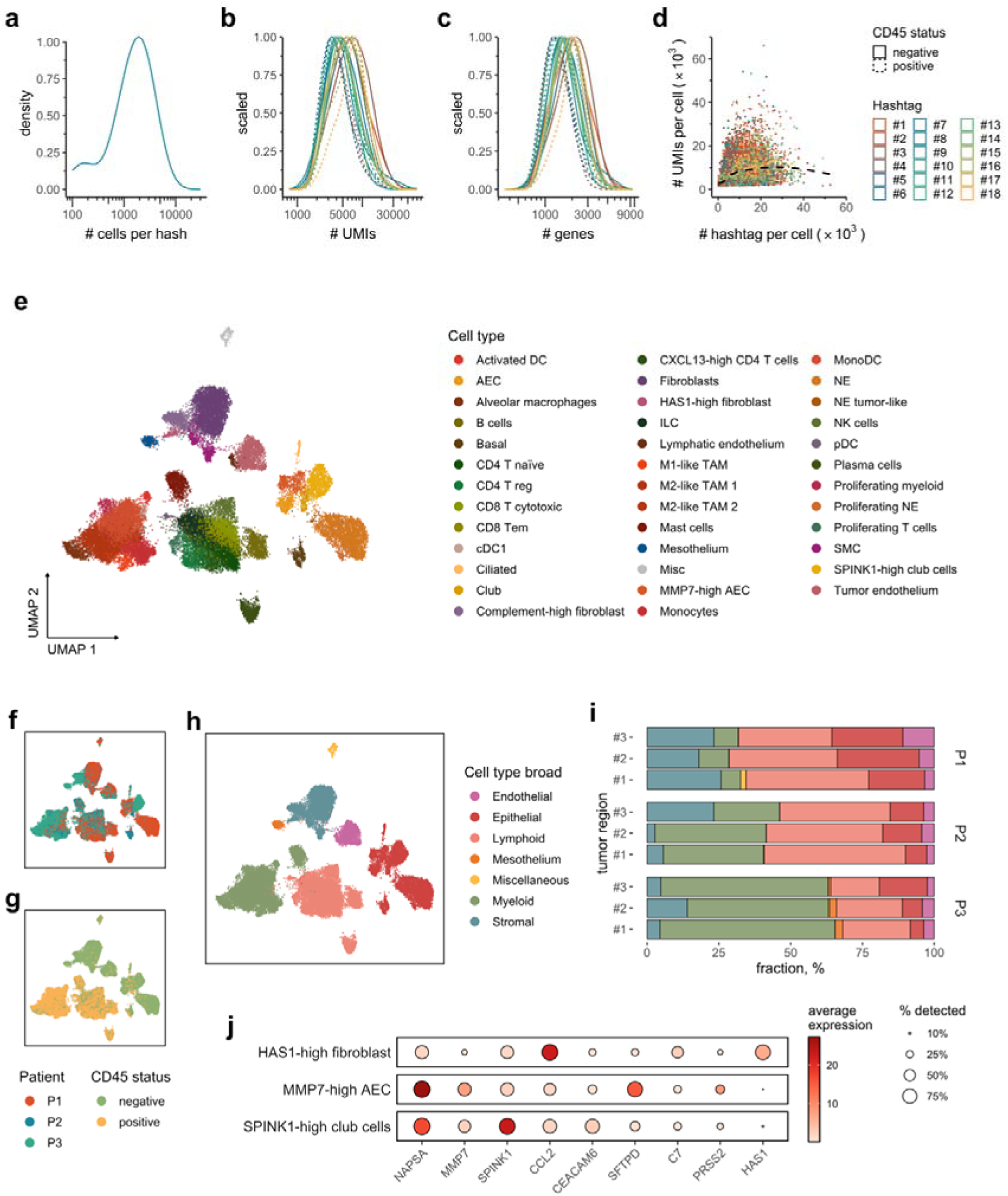
scRNA-seq of methanol-fixed and hashtag-indexed lung carcinoma cells. The plots display the probability distributions of **a)** cell number, **b)** total UMI count, and **c)** number of detected genes per individual hashtag, delineated by CD45 status. **d)** Relationship between UMI and hashtag counts per cell, where x-axis displays hashtag counts and y-axis UMI counts of the same cell barcode (dot coloured by an assigned hashtag). The black dashed curve is fitted on the data using loess method and shows a decrease in UMI count, when more reads are mapped to hashtags. **e**) An annotated UMAP of all lung carcinoma cells (n=32’937) displays high heterogeneity of cellular phenotypes in the tumor microenvironment. **f)** and **g)** UMAP colored by patient ID and CD45 status, respectively. **h)** UMAP representation colored by broad cell type categories. **i)** Sample composition analysis by broad cell type shows inter-patient compositional variability. **j)** Marker gene expression in selected cell populations, displayed by CP10k-normalized mean expression (represented by color) and fraction of cells expressing those genes (indicated by dot size). AEC – alveolar epithelial cells, cDC1 – conventional dendritic cells type I, ILC – innate lymphoid cells, NE – neuroendocrine cells, pDC – plasmacytoid dendritic cells, SMC – smooth muscle cells, TAM – tumor associated macrophages.

Other non-immune cells in our lung carcinoma atlas included lymphatic (*CCL21, NR2F2*) and tumor endothelial cells (*CLDN5, CLEC14A*), mesothelium (*MSLN, UPK3B*), smooth muscle cells (SMC, *ACTA2, TAGLN*) and diverse group of fibroblasts (*COL1A2, FN1*) which included two transcriptionally distinct groups involved in inflammation (Fig. 5e). Complement-high fibroblasts had an unusually high expression of complement system constituents (i.e. *C7, C3, CFD*) indicative of inflammatory processes in the tumor microenvironment (Supplementary Fig. 5b). Another fibroblast population was enriched for *HAS1* (Fig. 5j), similarly to an invasive fibroblast population recently discovered in fibrotic lungs [65]. Moreover, this population upregulated a potent chemokine for monocytes, *CCL2,* and other inflammatory factors, such as cytokines *CXCL1, CXCL2* and *IL6* (Supplementary Fig. 5b). These rare inflammatory fibroblast populations might be of interest for future detailed studies.

Consistent with previous reports [7, 59, 60] the myeloid compartment in our data comprised mast cells (*TPSB2, TPSAB1*), monocytes (*S100A9, FCN1*), conventional type 1 dendritic cells (*CLEC9A, CST3*), activated dendritic cells (*CCR7, CCL22*), monocyte-like dendritic cells (*CCL17, CLEC10A*), alveolar macrophages (*MARCO, FABP4*) as well as M1– and M2– like subpopulations of tumor-associated macrophages (TAM). The lymphoid compartment consisted of innate lymphoid cells (ILC, expressed *CD3D*), B cells (*CD79A, MS4A1*), plasma cells (*IGHG4, JCHAIN*), plasmacytoid dendritic cells (*LILRA4, CLIC3*), NK cells (*NKG7, GZMB*) and a large group of diverse T cell phenotypes. Specifically, within the T cell population, we captured CD4 regulatory T cells (*FOXP3, CTLA4*), naïve CD4 T cells (*IL7R, CCR7*), effector memory CD8 T cells (*CD52, S100A4*), cytotoxic CD8 T cells (*GZMA, CCL4*) and CXCL13-high CD4 T cells (Fig. 5e). Interestingly, the tumor samples comprising CXCL13-high T cell phenotype coincided with high count of B cells, therefore, supporting recent findings that CXCL13 acts as a potent attractant for B and other immune cells [66] (Supplementary Fig. 5c). Moreover, it was reported that the abundance of PDCD1-high CXCL13 producing CD8 T cells predicts response to PD-1 blockade therapy and correlates with increased overall survival in non-small cell lung cancer [67]. Overall, these findings clearly illustrate the power of inDrops-2 to recover clinically relevant cell phenotypes from biospecimens that underwent preservation, long-term storage and multiplexing, to obtain gene expression profiles of tens of thousands of single-cells in an efficient and inexpensive manner.

## Discussion

Advancements in high-throughput single-cell RNA-seq technologies [1] and computational methods [68] have opened new possibilities for investigating the gene expression programs and cellular composition in both normal and pathological conditions at unprecedented resolution and scale. As the range of scRNA-seq applications continues to expand across different domains of biomedical and biological sciences [16, 17, 69], there is a constant need for systems that not only deliver high throughput and sensitivity, but are also cost-effective. This is particularly relevant for analysis of biospecimens characterized by high heterogeneity, such as human tissues or cancer, necessitating the profiling of a large number of cells. Moreover, diverse needs of researchers often require versatile scRNA-seq platforms that can accommodate a broad range of samples and workflows.

Here we present an open-source scRNA-seq platform, inDrops-2, which enables high-throughput single cell transcriptomic studies with the transcript and gene detection matching that of state-of-the-art commercial platforms (i.e. 10X genomics Chromium v3), yet at the 6-fold lower cost (Supplementary Table S4). The system is highly flexible and customizable, providing a straightforward option to implement user-specific workflows. For instance, we implemented two popular scRNA-seq protocols; one based on linear amplification of cDNA using *in vitro* transcription and named as inDrops-2 (IVT), and another based on template switching reaction followed by exponential cDNA amplification, or inDrops-2 (TS). While both techniques were found to be well suited for identifying different cell types and detected similar transcript count, yet they also showed important differences. The scRNA-seq workflow based on exponential cDNA amplification following TS-reaction shows lower technical variability and requires less labor to obtain sequencing results. However, due to limited efficiency of template switching reaction, the scRNA-seq libraries tend to be enriched in shorter (≤14 kb) genes, including a significant fraction of ribosomal proteins. Given the median length size for protein-coding genes in humans to be 26 kb [70], the aforementioned technical bias might skew conclusions in certain biological contexts. For example, it has been reported that longer genes tend to be associated with cell development, complex diseases and cancer, while short genes are common to biological processes that require fast response such as the immune system [71]. There are indications that transcript length also plays a role in aging [72]. Therefore, the scRNA-seq libraries constructed following linear amplification of cDNA by *in vitro* transcription, although being labor intensive and relying on advanced molecular biology skillset, exhibit higher complexity and capture more genes per single cell, including non-coding RNAs, thus making it a potentially better choice for studying gene expression at a whole genome level.

In addition to improving scRNA-seq performance on live cells, we implemented a cell preservation procedure that safeguards intracellular mRNA from degradation and alleviates the technical challenges of working with primary cells that are prone to uncontrollable transcriptional changes during handling. Importantly, while our procedure is loosely built on previous reports [49–51], yet in contrast to others we explore gentle cell rehydration process to ensure minimal cell clumping and loss, making it applicable to clinical samples of limited availability (n ≤ 50,000 cells). The data quality of rehydrated cells was high, and matched current standards in the field, with UMI/gene capture similar to live cells using commercial platforms (10X Chromium V3). Taking one step further we explored chemical hashing (indexing) of dehydrated cells based on Dies-Alder reaction [52]. As a proof-of-concept we performed multiregional profiling of lung carcinomas by hash-tagging 18 samples that were methanol-preserved after acquiring them from several patients. Not only we captured all major specialized lung epithelial, infiltrating stromal and immune cell phenotypes [7, 59, 60], but also identified cells with potential significance in immunotherapy, such as the CXCL13 producing CD4 T cells. These results underscore the broad potential of inDrops-2 in biomedical research, especially in scenarios where characterization of complex diseases at single-cell level are of high relevance. As a side note, chemical cell hashing with DNA oligonucleotides (ClickTags) of methanol-fixed cells not only minimizes technical batch effects, but also benefits data analysis by facilitating unbiased removal of cell doublets all while preserving the clinically relevant cellular phenotypes.

Being an open and flexible platform inDrops-2 can accommodate diverse workflows matching user-specific needs and has a potential to further democratize single cell technologies. Indeed, it has recently been shown that inDrops-based platforms such as VASA-Seq [35] outperforms commercial and plate-based methods in terms of gene and transcript capture, while spinDrops [73] benefits applications based on target cell enrichment by on-chip sorting. Beyond transcriptomics, inDrops-based platforms have been successfully tailored to probe other –omic modalities in individual cells such as open chromatin by HyDrop [33] or genome by Microbe-Seq [74], to name a few. To conclude, inDrops-2 represents an affordable, sensitive, and adjustable open-source system that could expand the scope of single cell –omic applications, and further enhance the scalability of scRNA-Seq experiments.

## DATA AND CODE AVAILABILITY

Single-cell RNA-seq data presented in this work have been deposited at the European Nucleotide Archive (ENA) under the accession number PRJEB71611. Code used to process data and generate the main results underlying the publication is available at https://github.com/mazutislab/indrops-2 repository.

## ACKNOWLEDGEMENTS

This work received funding from European Regional Development Fund [01.2.2-LMT-K-718-04-0002] under grant agreement with the Research Council of Lithuania. Part of this work was also funded by The Alan and Sandra Gerry Metastasis and Tumor Ecosystems Center. S.J. was supported by the European Union’s Horizon 2020 research and innovation programme under the Marie Skłodowska-Curie grant agreement no. 101030265. We are grateful to Ignas Masilionis for wet-lab assistance, and the members of the SCRI at the Sloan Kettering Institute for their valuable support and kind assistance.

## AUTHOR CONTRIBUTIONS

SJ, KG, VK and JZ data analysis and interpretation; KG, VK and JN method development, single-cell RNA-seq experiments; AQV biospecimen acquisition, logistics and processing; L.M. study design, supervision and funding acquisition. LM wrote the initial draft of the manuscript; SJ, KG, JZ and LM revised the manuscript. All authors have read and approved the final manuscript.

## Materials and Methods

### Cell lines

Cryopreserved K-562 cells (ATCC, CCl-243) were stored in vapor phase nitrogen until use. Murine KP lung adenocarcinoma cell line [46] was a kind gift by Dr. Stella Paffenholz (MSKCC). Cell culture was maintained in 25 cm^2^ culture flask in 5 ml volume of Iscove’s Modified Dulbecco’s Medium, IMDM (Gibco, 31980030) supplemented with 10% fetal bovine serum, FBS (Gibco, 10270-106) and 1X penicillin-streptomycin, PS (Gibco, 15140122) under 5% CO_2_ and at 37 °C. Cells were harvested at ∼10^6^ cells/ml, collected into 15 ml conical tubes, pelleted at 300*g* for 5 min and washed twice in ice-cold 1x DPBS with 0.05% (w/v) BSA. The cell count and viability were quantified using the Countess II cell counter and 0.2% trypan blue staining.

### Human PBMC and bone marrow derived CD34+ stem/progenitor cells

Cryopreserved primary peripheral blood mononuclear cells (PBMC) from healthy donors were purchased from ATCC (PCS-800-011), while bone marrow stem/progenitor CD34+ cells from healthy donors were purchased from AllCells, LLC. (ABM022F) and stored in vapor phase nitrogen until use. Prior to scRNA-seq, a vial with frozen cells was removed from the liquid nitrogen tank and thawed at 37 °C in a water bath for 2-3 min. Next, the vial content (∼1 ml) was transferred to a 50-ml conical tube and slowly diluted with 1 ml of warm (∼37 °C) cell culture medium (IMDM with 10% FBS). To prevent osmolysis, warm medium was added dropwise while gently rocking the 50-ml tube with a hand. The thawed cells were serially diluted in 5-steps with 1:1 volume addition of warm medium and 2-min incubation between each step, until a final 32-ml volume was reached. The cell suspension was then pelleted at 300*g* for 5 min in a swinging bucket centrifuge. Supernatant was discarded and the cell pellet was washed twice in ice-cold 1x DPBS with 0.05% (w/v) BSA. The cell count and viability were determined with Countess II cell counter.

### Human lung cancer biospecimen acquisition and processing

The patients with lung adenocarcinoma (LUAD) or small cell lung carcinoma (SCLC) undergoing a surgical resection at Memorial Sloan Kettering Cancer Center (MSKCC) provided informed consent through an Institutional Review Board-approved biospecimen collection and analysis protocol. Each specimen was cut into three pieces, approx. 5-10 mm^3^ in size, and processed according to the dissociation protocol reported in the past [57]. Each sample was dissociated for 15 min at 37 °C on the GentleMACS Octo Dissociator with Heaters (Miltenyi) using Human Tumor Dissociation Kit (Miltenyi Biotec). Following tissue dissociation, the cell suspension was passed through 35 µm Cell Strainer Snap Cap and treated with red blood lysis (ACK buffer, Lonza) for 2 min at room temp. One LUAD sample was resuspended in PBS with 0.04% BSA and processed fresh for single cell encapsulation, while for the rest of the specimens, the cells were stained with live dye (Calcein AM) and PE anti-human CD45 antibody (BioLegend, cat no 368510) mixture, and using BD FACS Aria II instrument sorted into CD45+ and CD45-compartments. The sorted cells were spun down for 5 min at 300*g* in a swinging bucket centrifuge at 4 °C, and resuspended in 90% methanol. The methanol-preserved cells were transferred to –80 °C until all specimens were acquired.

### Methanol-based cell preservation

Cell preservation in methanol was adopted following the reports by Alles et al., 2017 and Chen et al., 2018, with some modifications. Specifically, the live cells were first transferred to DNA LoBind tube (Eppendorf, 0030108051), pelleted at 300*g* for 5 min at 4 °C and gently resuspended in 100 μl ice-cold 1x DPBS. Next, 900 μl of ice-cold methanol was slowly added in a dropwise manner while gently rocking the tube; this prevents cells from clumping and osmolysis. Once suspended in 90% methanol, the cells were incubated on ice for another 15 min and then transferred to –20 °C or –80 °C for a long-term storage. Refer to the Supplementary Protocol 3 for a comprehensive step-by-step procedure.

### Rehydration of methanol-preserved cells

The Supplementary Protocol 3 provides detailed description for rehydrating the methanol-fixed cells. Briefly, the tube with cells preserved in methanol was placed on ice for 15 min and then centrifuged at 1000*g* for 10 min in a swinging bucket centrifuge set at 4 °C. Most of the supernatant was removed leaving ∼50 µl on top of the cell pellet. Next, the cell pellet was resuspended in 400 μl of ice-cold Rehydration Buffer 1 (3X SSC, 80 mM DTT, 0.2% BSA, 1 U/µl RNase Inhibitor) and entire suspension transferred onto a centrifugal tube filter (Millipore, UFC30DV25), that was earlier pretreated with 1% BSA. The column was centrifuged at 50*g* for 45 s using 4 °C centrifuge. The flow through fraction was discarded. The cell suspension that was retained on top of the filter (∼50 μl volume) was washed two more times with an ice-cold Rehydration Buffer 1 and once with an ice-cold Rehydration Buffer 2 (1X SSC, 40 mM DTT, 0.1% BSA, 1 U/µl RNase Inhibitor). After final wash, the rehydrated cells were retrieved from the filter membrane, counted under hemocytometer and processed on 10x Chromium or inDrops-2 (TS) platform.

### scRNA-Seq using 10x Chromium platform

Single cell encapsulation and mRNA barcoding on 10X Genomics Chromium instrument was performed with Single Cell 3’ Library and Gel Bead Kit V3 reagent kit, following the vendor’s manual (CG00183 rev B). Briefly, for each sample a suspension of cells (viability 70-95%) were loaded onto Chromium microfluidics chip targeting for recovery of ∼5,000 single-cells with 3.9% multiplet rate. For cells resuspended in rehydration buffer 2 (1X SSC, 40 mM DTT, 0.1% BSA, 1 U/µl RNase Inhibitor), the cells were first concentrated by centrifugation to ∼2000 cells/µl and 4 µl were mixed with the corresponding V3 reagents as an input for encapsulation. Following the reverse transcription, the emulsion droplets were broken and barcoded cDNA purified with Dynabeads, followed by 12 cycles of PCR: 98 °C for 180 s, 12-cycles (98 °C for 15 s, 67 °C for 20 s, 72 °C for 60 s), and 72 °C for 60 s. The PCR-amplified barcoded cDNA was diluted to 50 ng, fragmented with the reagents provided in the kit, purified with SPRIselect beads (Beckman Coulter, B23318), and ligated to the sequencing adapters. The ligation product was amplified by PCR: 98 °C for 45 s, 14-cycles (98 °C for 20 s, 54 °C for 30 s, 72 °C for 20 s), and 72 °C for 60 s. The final DNA library was double-size purified (0.6-0.8×) with SPRI beads and sequenced on the Illumina NovaSeq 6000 platform (R1 – 26 cycles; i7 – 8 cycles; R2 – 70 or more cycles) at a depth of 10’000-50’000 reads per cell.

### Single cell encapsulation and mRNA barcoding using inDrops platform

Single cell suspensions (∼400 cells/µl) were prepared in 1x DPBS supplemented with 0.04% (w/v) BSA and 16% OptiPrep (M1248-100, BioVision). The OptiPrep is used to increase the density (ρ) of 1x PBS buffer to ρ*_sol_* = 1.044 g/ml, thereby suppressing cell sedimentation. Single cells suspensions were loaded onto microfluidics chip (Supplementary Fig. 1) along with barcoded hydrogel beads (either V1 or V2 design, see Supplementary Table S1) and 2x RT-lysis mixture (see below). The barcoded hydrogel beads were suspended in 1x First Strand buffer (TFS, 18080044) supplemented with 0.3% (v/v) IGEPAL CA-630 (Sigma-Aldrich, 18896-50ML). The 2x RT-lysis mixture for inDrops-1 protocol comprised: 24 U/µl SuperScript III Reverse Transcriptase (TFS, 18080044), 1.3 U/µl SUPERase-In (TFS, AM2696), 0.83x First Strand buffer (TFS, 18080044), 0.6% (v/v) IGEPAL CA-630 (Sigma-Aldrich, 18896-50ML), 5 mM DTT (TFS, 00561515), 11 mM MgCl_2_ (Ambion, AM9530G), 65 mM Tris-HCl (Invitrogen, 15568-025) and 1 mM dNTPs (TFS, R0192). The 2x RT-lysis mixture for inDrops-2 (IVT) protocol comprised: 24U/µl of Maxima H minus (TFS, EP0751), 2U/µl of RiboLock RNase inhibitor (TFS, EO0381), 2x RT buffer (TFS, EP0752), 1 mM dNTP, 0.6% (v/v) IGEPAL CA-630. The 2x RT-lysis mix for inDrops-2 (TS) protocol comprised: 24U/µl of Maxima H minus, 2U/µl of RiboLock RNase inhibitor, 2x RT buffer, 1 mM dNTP, 0.6% (v/v) IGEPAL CA-630 and 50 μM template switching oligonucleotide, TSO (Supplementary Table S1). When testing the performance of Super Script IV enzyme, the Maxima H minus enzyme and RT buffer was replaced with 24 U/µl SS-IV enzyme and corresponding SS-IV buffer (Invitrogen, 18090010), while keeping other ingredients in the reaction mixture the same. The microfluidics platform was operated under two flow regimes; standard and high-throughput. For a standard run, the flow rates were set at 250, 250 and 50-70 µl/hr for cells (diluted at ∼500 cells/µl), 2x RT-lysis mixture and barcoded hydrogel beads, respectively. The droplet stabilization oil was set at 550 µl/hr. For the high-throughput run, the flow rates were set at 100, 900, 150 and 1200 µl/hr for cells (diluted at ∼2000 cells/µl), RT-lysis mixture, barcoded beads and carrier oil, respectively. Emulsion droplets were collected on-ice for 20 to 60 min. Next, the barcoded RT primers were photo-released by exposing the tube with an emulsion to a 350-nm light either using LED device (Atrandi, MHT-LAS1) for 20 seconds, or UVP lamp (UVP, cat. no. 95-0127-01) for 5 min. The emulsion was then transferred onto a heat block to initiate cDNA synthesis at either 42 °C or 50L°C for 60 min followed by the heat inactivation at 75 °C for 15 min (for inDrops-1) or at 85 °C for 5 min (for inDrops-2), or otherwise as indicated. The post-RT droplets were aliquoted into separate tubes (each containing ∼20.000 droplets) and then broken by adding 1H,1H,2H,2H-perfluorooctanol up to 10% (v/v). After a quick spin for 30 sec at 300*g*, the supernatant was transferred onto filter column (Zymo, C1004-250). The flow-through fraction containing barcoded-cDNA was collected into a new 1.5 ml DNA LoBind tube (Eppendorf, 0030108051) by centrifugation for 1 min at 1000*g*. After this step, the barcoded-cDNA can be stored at 4 °C overnight, or processed further to construct the sequencing library as indicated below.

### inDrops-1 sequencing library preparation

The barcoded-cDNA was diluted to 80 µl with nuclease-free water and treated with an enzyme cocktail comprising exonuclease I, restriction endonuclease HinfI and alkaline phosphatase. Specifically, the inDrops-v1 libraries were digested with 40U of ExoI (TFS, EN0581), 4 µl of FastDigest HinFI (TFS, FD0804) and 1U of FastAP (TFS, EF0654) for 15 min at 37 °C and purified using 1.2× SPRIselect beads (Beckman Coulter, B23318). The inDrops-v1.1 libraries were digested with 40U of ExoI and 1U of FastAP for 15 min at 37 °C, and purified using 1.2× SPRIselect beads. The inDrops-v1.2 iteration excluded enzymatic treatment and instead the libraries were purified with 0.8× SPRIselect beads. The purified cDNA was converted to double stranded DNA (dsDNA) by performing second strand synthesis with NEBNext Ultra II Non-Directional RNA Second Strand Synthesis Module (NEB, E6112) in 20 µl reaction volume for 2.5 h at 16 °C and 20 min at 65 °C. The dsDNA was then linearly amplified using HiScribe T7 High Yield RNA Synthesis Kit (NEB, E2040S) for 15 h at 37 °C. Reaction products, in the form of RNA, were purified using 1.2× SPRIselect beads and their quality as well as yield was evaluated with RNA Pico Assay on Agilent Bioanalyzer 2100 instrument. The amplified material was fragmented with RNA fragmentation reagent (Ambion, AM8740) at 70 °C for 2.5 min, terminated with 20 mM EDTA and purified with 1.2× SPRIselect beads. After purification, fragmented RNA library was mixed with 5 µM of PE2-N6 primer (Supplementary Table S1) and reverse transcribed using PrimeScript RTase (Takara, SD0418) for 60 min at 42 °C. The resulting cDNA library was purified with 1.2× SPRIselect beads and amplified by 12-cycles of PCR: 98 °C for 2 min, 2-cycles (98 °C for 20 s, 55 °C for 30 s, 72 °C for 40 s), 10-cycles (98 °C for 20 s, 65 °C for 30 s, 72 °C for 40 s), and 72 °C for 5 min. DNA amplification was conducted with Kapa HiFi HotStart PCR mix (Kapa Biosystems) using PE1 and PE2 indexing primers (Supplementary Table S1). The amplified and indexed libraries were further purified using SPRIselect dual-size selection (0.6-0.8×). The library size was evaluated using HS DNA assay (Agilent Technologies, 5067-4626). The libraries were adjusted to 10 pM spiked in with 15% PhiX Control v3 Library and sequenced on the NextSeq550 and HiSeq2500 (Illumina) platform (R1 – 54 cycles; i7 – 8 cycles; R2 – 35 cycles or more) at a depth of ∼20’000 reads per cell.

### inDrops-2 (IVT) sequencing library preparation

The detailed protocol for constructing inDrops-2 (IVT) libraries is provided as Supplementary Protocol 1. The barcoded-cDNA was diluted to 80 µl and purified using 0.8× SPRIselect beads (B23318, Beckman Coulter), and subjected to second-strand synthesis using NEBNext Ultra II Non-Directional RNA Second Strand Synthesis Module in 20 µl reaction volume for 2.5 h at 16 °C and 20 min at 65 °C. The second-strand synthesis reaction product was then linearly amplified using HiScribe T7 High Yield RNA Synthesis Kit for 15Lh at 37 °C. The reaction products, in the form of RNA, were purified using 0.8× SPRIselect beads and their quality as well as yield was evaluated using RNA Pico Assay (Agilent, 5067-1513). The RNA concentration was estimated based on UV absorption at 260 nm (NanoDrop) and diluted to 1000 ng/µl. Next, 9 µl of amplified RNA was fragmented with 1 µl RNA fragmentation reagent (Ambion, AM8740) at 70 °C for 1.5 min, and purified with ice-cold STOP mixture (9 µL of 10 mM Tris [pH 7.0], 25 µl of SPRIselect beads, 2 µl of 200 mM EDTA). The purified RNA was mixed with 5 µM of PE2-N6 primer (Supplementary Table S1), heated to 70 °C for 2 min and allowed to hybridize for 3 min on ice. The RNA was reverse transcribed using Maxima H minus enzyme at 30 °C for 10 min followed by 60 min at 42 °C and heat inactivation for 15 min at 70 °C. The cDNA was purified with 1.0× SPRIselect beads and amplified by Kapa HiFi HotStart PCR mix using PE1 and PE2 Illumina index primers (Supplementary Table S1). The PCR was set for 10-cycles: 98 °C for 2 min, 2-cycles (98 °C for 20 s, 55 °C for 30 s, 72 °C for 40 s), 8-cycles (98 °C for 20 s, 65 °C for 30 s, 72 °C for 40 s), and 72 °C for 5 min. The amplified libraries were purified using SPRIselect dual-size selection (0.6-0.8×), inspected on a High Sensitivity DNA Chip (Agilent Technologies, 5067-4626), and library yield quantified with Qubit dsDNA HS Assay kit. The final inDrops-2 (IVT) libraries were diluted to 10 pM, mixed with 7.5% PhiX Control v3 Library, and sequenced on the NextSeq550 and HiSeq2500 (Illumina) instrument (R1 – 54 cycles; i7 – 8 cycles, R2 – 35 cycles or more), at a depth of 10’000 to 50’000 reads per cell.

### inDrop-2 (TS) sequencing library preparation

The detailed protocol for constructing inDrops-2 (TS) libraries is provided as Supplementary Protocol 2. The barcoded-cDNA was purified with 0.8× SPRIselect beads and subjected to PCR (2× KAPA HiFi HotStart Ready mix, KK2500) using cDNA amplification primers (Supplementary Table S1) at 0.5 µM, and following PCR program: 98 °C for 3 min, 14-cycles [98 °C for 15 s, 67 °C for 20 s, 72 °C for 60 s], 72 °C for 1 min. The amplified DNA was purified with 0.8× SPRIselect beads and assessed with DNA HS assay on Agilent Bioanalyzer 2100. Next, the library for sequencing was constructed with FS DNA Library Prep Kit (NEB, E7805) by following the manual instructions. Specifically, 50 ng of amplified cDNA was fragmentated for 8 min at 37 °C, followed by end-repair and dA tailing for 30 min at 65°C. Then, the modified double-stranded ligation adapter (Supplementary Table S1) supplied at 55 nM was ligated to fragmented DNA for 15 min at 20 °C. The ligation product was purified using 0.8× SPRIselect beads, eluted in 40 µl nuclease-free water, and half of the material was subjected to amplification by PCR (NEBNext Ultra II Q5 Master Mix) using indexing primers (Supplementary Table S1) at 0.5 μM. The PCR was set for 14-cycles: 98 °C for 45 s, 14-cycles [98 °C for 20 s, 54 °C for 30 s, 72 °C for 20 s] and 72 °C for 1 min. The amplified DNA was purified using SPRIselect dual-size selection (0.6-0.8×) and the library size distribution was evaluated with DNA HS assay on Agilent Bioanalyzer 2100. The inDrop-2 (TS) libraries were sequenced on the MiSeq, HiSeq2500, NextSeq550 and NovaSeq6000 (Illumina) platforms, without PhiX spike-in. The sequencing parameters were R1 – 28 cycles; i7 – 8 cycles, R2 – between 35 and 92 cycles (depending on instrument and sequencing reagent kit), at a depth of >10’000-50’000 reads per cell.

### Cell hashing using click-chemistry oligonucleotide tags

Cell hashing with DNA oligonucleotides was adapted from the work by Gehring et al., 2020 [52] with following modifications:

1. ClickTag preparation. The 3’-amino modified oligos (Supplementary Table S5) were activated by 1 mM NHS-methyltetrazine (Click Chemistry Tools) in 50% DMSO for 60 min at 21 °C. The activated ClickTags were precipitated by ethanol and resuspended in 10 mM HEPES [pH 7.2] to yield the final concentration of 40 μM. Activated ClickTags were stored in the dark at –20 °C for up to 3 days, before use.
2. Cell hashing with ClickTags. To hashtag the cells with ClickTags, 25 µl of methanol-preserved cells (∼30’000 cells in total) were mixed with 1 mM NHS-TCO (Click Chemistry Tools) and incubated in the dark at 21 °C for 5 minutes. Next, the cell suspension was mixed with 3 µl of 40 µM activated hashtag oligonucleotide and incubated for 30 min on a rotating platform at room temperature. The reaction was terminated by quenching with a 10 mM Tris-HCl [pH 8.0] buffer supplemented with 50 µM methyltetrazine-DBCO (Click Chemistry Tools). The methanol-preserved cells conjugated with ClickTags were rehydrated following Supplementary Protocol 3.
3. scRNA-Seq of hash-tagged cells. The rehydrated hash-tagged cells were resuspended in a rehydration buffer (1X SSC, 40 mM DTT, 0.1% BSA, 1U/µl RiboLock RNase Inhibitor) with 20% Optiprep at a dilution of ∼2000 cells/µl. Then, cell suspension was loaded onto inDrops-2 microfluidics chip along with 1.1x RT-lysis mixture and barcoded hydrogel beads. The flow rates of the microfluidics platform were adjusted to compensate for the inhibitory effect of citric acid that is present in the rehydration buffer. As such, the flow rates were set at 100 µl/hr for cells, 900 µl/hr for 1.1x RT-lysis mixture and 100-150 µl/hr for barcoded hydrogel beads, respectively. The droplet stabilization oil was set at 1200 µl/hr. The emulsion was collected on-ice for 20 min and after release of barcoded RT primers the cDNA synthesis was initiated at 42 °C for 90 min followed by 85 °C for 5 min. The post-RT droplets were broken with 10% perfluorooctanol and supernatant containing barcoded-cDNA was collected by passing through a filter column (Zymo, C1004-250) at 1000*g* for 1 min.
4. ClickTag sequencing library preparation. Following inDrops-2 (TS) approach the cDNA derived from hashtags and mRNA was co-purified using 2× AMPure beads. Then, purified material was amplified by 14-cycles of PCR (98 °C for 3 min, 14-cycles [98 °C for 15 s, 67 °C for 20 s, 72 °C for 60 s], 72 °C for 1 min) using 0.5 µM of cDNA amplification oligos (Supplementary Table S1) that targeted mRNA-derived cDNA, and 2 µM of Hash_fwd_oligo (Supplementary Table S5) that targeted hashtag-derived cDNA. The amplified material was mixed with 0.6× volume of SPRIselect beads and eluted in two fractions. The magnetic bead bound fraction comprising mRNA-derived cDNA was processed following the inDrops-2 (TS) protocol (Supplementary Protocol 2). The supernatant fraction comprising hashtag-derived cDNA was mixed with SPRI beads to reach a final SPRI ratio of 1.6×, incubated at room temperature for 5 min, washed twice with 80% ethanol and eluted in 20 Lμl of nuclease-free water. The hashtag libraries were spiked with mRNA-derived cDNA library at a ratio of 1:10, and sequenced on the NovaSeq6000 (Illumina) platform (R1 – 28 cycles; i7 – 8 cycles, R2 – 70 cycles or more), at a depth of >10’000 reads per cell.

### RNA extraction and evaluation

Total RNA extraction was performed with TRIzol reagent (TFS, 15596026) following manufacturer’s instructions. The extracted RNA was further purified with RNA clean and Concentrator kit (Zymo Research, R1060). The RNA integrity number (RIN) was estimated using total RNA Pico Assay (Agilent Technologies, 5067-1513) on the Agilent Bioanalyzer 2100 instrument.

### Optimization of cDNA synthesis and purification

To arrive at optimal reaction conditions that can consistently generate high cDNA yields, the effect of various cell lysis agents, the RT enzymes and temperature, TSO concentration, and cDNA purification strategies were first tested in a bulk format (Supplementary Fig. 2). The RT reaction was performed in 20 µl and comprised a total of ∼20.000 K562 cells including 1X RT buffer, 0.5 µM RT primer (5’-CTACACGACGCTCTTCCGATCTNNNNNNTTTTTTTTTTTTTTTTTTTTTTTTTTTTTT), 0.5 mM dNTP, 1 U/µl RiboLock RNase inhibitor, 0.1% (v/v) of corresponding lysis agent, varying amount of TSO, varying amount and type of RT enzyme. The reaction was carried out at 42 °C for 60 min followed by 5 min at 85 °C. The lysis agents tested in this work included IGEPAL CA-630 (Sigma-Aldrich, 18896-50ML), Tween 20 (Sigma-Aldrich, P9416-100mL), Triton X-100 (Sigma-Aldrich, 93426-100mL), Brij58 (Thermo Scientific, 28336), Digitonin (Invitrogen, BN2006), n-octyl-β-glucopyranoside (Roth, CN23.1), all at 0.1% (w/v). The impact of TSO amount was evaluated in the range of 5 to 50 µM (Supplementary Fig. 2c). The RT enzymes tested included Maxima H minus (TFS, EP0751) and SuperScript IV (Invitrogen, 18090050). In all cases, the cDNA was purified with 1.2× SPRIselect beads except when evaluating different purification strategies, which included; 1.2× SPRIselect beads, Zymo DNA Clean & Concentrator kit (Zymo Research, R1060), GndHCl buffer (7.3 M guanidinium chloride, 100 mM Tris-HCl [pH 6.2]) and GndSCN buffer (3 M guanidinium isothiocyanate, 33% isopropanol, 4% Triton X-100, 20 mM Tris-HCl pH [6.2]) (Supplementary Fig. 2e) [75]. cDNA purification with SPRIselect beads and Zymo DNA Clean & Concentrator kit was performed according to manufacturer’s recommendations. To purify cDNA with in Dynabeads first, Dynabeads were mixed with either GndHCl or GndSCN buffers in 1:24 ratio. Then post-RT reaction mix was purified by adding 2.5× volumes of Dynabeads in binding buffer and incubating for 5 min at room temperature. After incubation tubes were placed on a magnetic stand for magnetic beads to settle. Supernatant was removed and the bead pallet was washed 2-times with 180 µl of 80% ethanol. After washing, the beads were left to dry for 2 min at room temperature. To elute cDNA, the bead pellet was resuspended in 20 µl nuclease-free water, incubated for 2 min at room temperature and the supernatant collected after settling the magnetic beads on a magnetic stand. Once purified, the cDNA was amplified by 14-cycles PCR (KAPA): 98 °C for 3 min, 14-cycles [98 °C for 15 s, 67 °C for 20 s, 72 °C for 60 s], 72 °C for 1 min, using 0.5 µM cDNA amplification primers (Supplementary Table S1). The amplified cDNA was purified with 1.2× SPRIselect beads, diluted 6-times in pure MQ-water and analyzed with DNA HS assay on Agilent Bioanalyzer 2100.

### Optimization of inDrops-2 (TS) library construction

To arrive at the final inDrops-2 (TS) library construction protocol, we tested multiple reaction conditions in bulk format. Specifically, for each condition tested we performed cDNA synthesis in 20 µl volume comprising ∼20.000 cells (K-562), 1X RT buffer, 0.1% (v/v) IGEPAL CA-630, 0.5 mM dNTP, 25 µM TSO, 0.5 µM RT primer (see RT-trim3 primer on Supplementary Table S1), 1 U/µl RiboLock RNase inhibitor and 10 U/µl Maxima H-minus RTase. The RT reaction was carried out at 42 °C for 60 min, terminated at 85 °C for 5 min and cDNA purified with 1.2× SPRIselect beads. Following 14-cycles PCR (KAPA) the amplified cDNA was purified with 1.2× SPRIselect beads, and 50 ng of material was fragmented for 8 min at 37 °C and A-tailed for 30 min at 65 °C using NEBNext Ultra II FS DNA Library Prep Kit (NEB, E7805S). The fragmented and dA-tailed DNA library was ligated to dsDNA adapter having both forward (Ligation FWD primer) and reverse primers (Ligation REV primer) blocked at 3’ and 5’ ends, respectively (Supplementary Table S1). The adapter ligation reaction was performed for 15 min at 20 °C using a Ligation Master Mix provided with NEBNext Ultra II FS DNA Library Prep Kit. The libraries were purified with 0.8× SPRIselect beads, PCR amplified and analyzed on a High Sensitivity DNA Chip. The results of these efforts are provided in Supplementary Fig. 2.

### Microfluidics platform setup

We used a custom-built microfluidics platform reported previously in details [38] as well as open-source microfluidics platform Onyx (Atrandi Biosciences) both of which provide experimental flexibility for encapsulating cells at a desirable throughput and reaction conditions. The microfluidic chips were made of the polydimethylsiloxane (PDMS) bound to a microscope glass slide, and having rectangular microchannels 80 µm height. When performing single-cell RNA-Seq, the samples were injected into the microfluidics chip via PTFE tubing (int. 0.56 mm; ext. 1.07 mm, Atrandi (MAN-TUB2)) connected to 1 mL syringes (Omnifix-F, BRAUN) and 0.6 x 25 mm Neolus needles (Terumo). The syringe and tubing for barcoded hydrogel beads was wrapped in aluminum foil to protect photo-cleavable primers from illumination by ambient light. The flow rates of liquids and carrier oil were controlled by syringe pumps (PHD 2000, Harvard Apparatus). Emulsions were collected off-chip into 1.5 ml DNA LoBind tubes (Eppendorf, 0030108051) placed on a cooling rack (4 °C).

### Barcoded hydrogel bead synthesis

Synthesis of V1 beads. The hydrogel beads carrying DNA barcode sets were synthesized following the previously described protocol [38] and employing the Agilent Bravo Automated Liquid Handling Platform. At first, the 58 ± 2 µm size acrylamide-based hydrogel beads carrying 50 µM of photo-cleavable DNA stub: 5’-/5Acryd/PC/CGATTGATCAACG TAATACGACTCACTATAGGGATACCATCTACACTCTTTCCCTACACGACGCTCTTCCG-3’, where 5Acryd is an acrydite moiety, PC is a photo-cleavable spacer) were generated using a microfluidics chip (Atrandi, MCN-G5) and stored at dark at 4 °C in a Washing Buffer (10 mM Tris-HCl [pH 8.0], 0.1 mM EDTA and 0.1% Tween-20) until further use. Next, the DNA barcodes were attached to the DNA stub on hydrogel beads by conducting two rounds of primer extension reaction in a combinatorial split-and-pool manner. In the first round the hydrogel beads were washed 2-times in isothermal buffer (20 mM Tris-HCl [pH 8.8], 10 mM (NH_4_)_2_SO_4_, 50 mM KCl, 2 mM MgSO_4_, 0.1% Tween 20) and 1-time in the isothermal buffer supplemented with 0.3 mM dNTP (each). After diluting hydrogel bead suspension to ∼10.000 beads/µl, the Agilent Bravo Liquid Handling Platform, was used to distribute the beads in the four 96-well plates (10 μl of beads per single well), preloaded with 5-µl of barcoded oligonucleotides (1^st^ set of sub-barcodes) at 50 µM concentration. (Supplementary Table S6). After heating the plates at 85 °C for 2 min and 60 °C for 1 hour, the 5-µl of Bst 2.0 (NEB, M0537S) enzyme mixture (1× isothermal buffer, 0.3 mM dNTP and 1.8U Bst 2.0 enzyme) was added to each well, gently mixed, and the primer extension reaction initiated at 60 °C. After 30 min of incubation the reaction was terminated by adding 25 µl of STOP solution (10 mM Tris-HCl [pH 8.0], 50 mM EDTA; 0.1% Tween 20 and 100 mM KCl). The beads were collected into a single 50 ml tube and the second strand was removed by alkaline denaturation. For that purpose, the beads were washed 5-times in Denaturation Solution (0.1 M NaOH, 0.5% Brij 35P) with 5 min incubations between the washes. The alkaline solution was neutralized with one volume of Neutralization Buffer (100 mM Tris-HCl [pH 8.0], 100 mM NaCl, 10 mM EDTA, 0.1% Tween 20), and then washed twice in a Washing Buffer until further use. At this step the hydrogel beads had 1^st^ set of sub-barcodes attached to them in a single-stranded form, thus making them suitable for the 2^nd^ round of barcoding. Following the same procedure as described above the hydrogel beads were washed twice in the isothermal buffer, aliquoted in four 96-well plates preloaded with 2^nd^ set of sub-barcodes (Supplementary Table S6), and then subjected to the second round of barcoding followed by dsDNA denaturation, neutralization and washing. To remove the ssDNA primers that lack poly(dT) tails the beads were resuspended in 1× ExoI buffer (TFS, EN0581), hybridized to 50 µM poly(dA)24 primer for 10 minutes and treated with Exonuclease I enzyme (TFS, EN0581) for 30 minutes at room temperature. Next, Klenow Exo-minus reaction mixture comprising 1× FD buffer (TFS, B64), 0.25U Klenow Exo minus fragment (TFS, EP0421) and 8.7 mM dNTPs was added to the hydrogel bead suspension at a dilution 1:10 (Klenow reaction mixture:hydrogel bead suspension), and incubated at 37 °C for 60 min. The hydrogel beads were then washed 6-times in Denaturation Solution with 3 min incubations between the washes and 2-times in Neutralization Buffer. The beads were filtered through a 70 μm strainer, resuspended in a Washing Buffer and stored at 4 °C until further use. The quality and yield of fully-barcoded DNA primers was evaluated with total RNA Pico Assay (Agilent Technologies, 5067-1513) and by fluorescent *in situ* hybridization (FISH) targeting 3’ poly(dT) tail, as described by Zilionis et al., 2017. The full-length primer sequence on V2 hydrogel beads was: 5’-/5Acryd/PC/CGATGACGTAATACGACTCACTATAGGGATACCACCATGG <u>CTCTTTCCCTACACGACGCTCTTCCGATCT</u>[12345678901]GAGTGATTGCTTGTGACGC CTT[12345678]NNNNNNNNTTTTTTTTTTTTTTTTTTT-3’, where 5Acryd is an acrydite moiety, PC is a photo-cleavable spacer, the letters in bold indicate T7 RNA promoter sequence, and underlined letters indicate the site for Illumina PE Read 1 Sequencing primer. The numbers indicate cell barcodes, which were designed to have 50% GC content and Hamming distance of ≥ 3 between each pair of barcodes. The barcode whitelist is provided in Supplementary Table S7.

Synthesis of V2 beads. The hydrogel beads 63 µm in size and carrying the photo-cleavable DNA stub (/5Acryd/CGATGACG(PC)CTACACGACGCTCTTC-3’, where Acryd is an acrydite moiety, PC is a photo-cleavable spacer) at 50 µM concentration were subjected to two rounds of combinatorial ligation reaction. The liquid handling operations were conducted using Agilent Bravo Liquid Handling Platform. At first, the 1^st^ set of sub-barcodes (Supplementary Table S8) were made double-stranded by mixing P_Bcd1 and RC_Bcd1 primers in equimolar amount, denaturating for 5 min at 95 °C and cooling to 4 °C at a rate 0.3 °C/min. The oligonucleotide duplexes (200 µM) were aliquoted into 4 x 96-well plates, 6.25-µl per well. Then, a master mix containing hydrogel beads carrying DNA stub and ligation mixture was distributed in the same 4 x 96-well plates such that each well would contain 12.5 μl of close-packed hydrogel beads,1.25 µl of 10× ligation buffer (NEB, B0202S), 0.3 μl of T4 DNA ligase I (NEB, M0202L), 6.25-µl of double-stranded barcoded oligonucleotides (1^st^ set of sub-barcodes) at 200 µM concentration and 4.7 µl of nuclease-free water. The ligation reaction was conducted overnight at room temperature while slowly rotating the plates on a vertical tube rotator (Biosan). The reaction was stopped by adding 50 μl of STOP buffer (50 mM Tris-HCl, [pH 8.0], 50 mM EDTA, 0.1% Tween 20) to each well, gently mixed by pipetting and all hydrogel beads pooled in a single 50-ml tube. After washing 5-times with 25 ml of 1× ligation buffer, the ligation reaction was conducted with a 2^nd^ set of barcodes. The hydrogel beads carrying 1^st^ set of barcodes were evenly distributed in four 96-well plates preloaded with 6.25µl of single-stranded barcoded oligonucleotides (2^nd^ set of sub-barcodes) at 200 µM concentration (Supplementary Table S8). The 25-µl volume reaction composition per single well comprised: 1× ligation buffer, ∼500.000 hydrogel beads, 50 µM bcd2 primer and 120 U of T4 DNA ligase I. The ligation reaction proceeded overnight at room temperature on a rotating platform to prevent hydrogel beads from settling. After completing two-step ligation, the beads were pooled and washed 5-times with 25-ml of Washing Buffer. To remove, the oligonucleotides lacking poly(dT) tails the hydrogel beads were resuspended in 1× Exo I buffer (TFS, EN0581), hybridized to 50 µM poly(dA)24 primer for 10 minutes and treated with Exonuclease I enzyme (TFS, EN0581) for 30 minutes at room temperature. The hydrogel beads were then washed 6-times in Denaturation Solution with 3 min incubations between the washes and 2-times in Neutralization Buffer. The beads were filtered through a 70 μm strainer, resuspended a Washing Buffer, and the quality as well as yield of fully-barcoded DNA primers was evaluated with total RNA Pico Assay (Agilent Technologies, 5067-1513) and by fluorescent *in situ* hybridization (FISH) targeting 3’ poly(dT) tail. The full-length V2 primer sequence on the beads was as follows: /5Acryd/CGATGACG/PC/<u>CTACACGACGCTCTTCCGATCT</u>[12345678]CATG[12345678]NN NNNNNNTTTTTTTTTTTTTTTTTT-3’, where 5Acryd is an acrydite moiety, PC is a photo-cleavable spacer, underlined letters indicate the site for Illumina P7 Read 1 Sequencing primer. The numbers indicate cell barcodes, which were designed to have 50% GC content and Hamming distance of ≥ 3 between each pair of barcodes. The barcode whitelist is provided in Supplementary Table S9.

### Single-cell RNA-seq data pre-processing

Each pair of the index 1 (i7) demultiplexed read 1 and 2 fastq files (containing cell barcode and transcript sequences, respectively) were processed using STAR v2.7.10a [76]. Specifically, the reads were mapped to the human GRCh38 genome (GENCODE v41) or the mouse GRCm38 genome (GENCODE M30) with default parameters using –-soloFeatures *GeneFull* to count genes (incl. exons and introns) and to generate cell ⨉ gene count matrices. To reduce the gene overlap-derived read loss (mapping to multiple features), read through transcripts and transcripts within pseudoautosomal regions were excluded from the GENCODE annotations and were limited to protein coding, lncRNA, IG/TR gene and pseudogene biotypes. Since only uniquely mapped genes were counted, for some libraries, transcript reads (read 2) were trimmed before the alignment to achieve an equal number of bases among compared libraries using seqkit v2.0.0 [77]. To define barcode geometry, barcode correction and UMI deduplication for different protocols, the parameters were the following: (i) for inDrops-1 and inDrops-2 libraries generated with barcodes v1: –-soloType *CB_UMI_Complex*, –-soloCBmatchWLtype *EditDist_2*, –-soloUMIdedup *Exact*, –-soloAdapterSequence *GAGTGATTGCTTGTGACGCCAA*, –-soloCBposition *0_0_2_-1 3_1_3_8*, –-soloUMIposition *3_9_3_16* (barcode list is provided in the Supplementary Table S7); (ii) for inDrops-2 (TS) with barcodes v2: –-soloType *CB_UMI_Complex*, –-soloCBmatchWLtype *EditDist_2*, –-soloUMIdedup *Exact*, –-soloCBposition *0_0_0_7 0_12_0_19*, –-soloUMIposition *0_20_0_27* (barcode list is provided in the Supplementary Table S9); (iii) for 10x Genomics v3: –-soloType *CB_UMI_Simple*, –-soloCBmatchWLtype *1MM_multi*, –-soloUMIdedup *1MM_CR*. Complete examples of the STAR parameters for each protocol are provided in the supplementary code (see Data and Code Availability). The processing of the methanol-fixed hashtag single-cell RNA-seq data was done using a combination of SEQC [78] and CITE-seq-Count [79] pipelines. The SEQC with default parameters for inDrops-2 (TS) (barcode version 2 (Supplementary Table S9) was used to obtain cell ⨉ gene count matrix, while the CITE-seq-Count (v1.4.4) with the default parameters and *--no_umi_correction* was used to count hashtag sequences (Supplementary Table S5) and to generate cell ⨉ hashtag count matrix. Cells were classified as singlets or doublets and assigned to corresponding samples with HashSolo [80].

### Single-cell RNA-seq data downsampling

The resulting bam files, tagged with corrected cell barcode (CB), UMI (UB) and gene (GN) information were parsed using the pysam v0.20.0 and down-sampled employing random sampling with the numpy v1.19.2 random.rand() approach. More explicitly, each cell barcode was either downsampled to a given number of raw reads (20’000 or 15’000) or to a given proportion of raw reads (5%, 10%, 20%, 30%, 40%, 50%, 60%, 70%, 80%, 90% and 100%), while counting the detected UMIs and unique genes. To generate saturation curves for each protocol version, the top quarter of cell barcodes based on UMI counts (n > 1000) were included. To evaluate whether the observed differences in UMI and gene count were statistically significant, two-tailed t-test from the rstatix v0.7.0 was applied in a pairwise manner on downsampled UMI and gene counts of cells that achieved required sequencing depth. The resulting P values were adjusted for multiple corrections by Benjamini-Hochberg false discovery rate (FDR). Custom scripts used for the down-sampling are provided in supplementary code (see Data and Code Availability).

### Quality control and doublet detection

To remove cell barcodes with low complexity, the cell x gene count matrices were filtered by UMI counts (>2300 UMIs for PBMC samples and >1500 UMIs for CD34^+^ bone marrow cells) and mitochondrial gene count fraction (>20% for all samples). For lung carcinoma samples cells were filtered based on hashtag data, removing cells that were assigned multiple hashtags. Doublets were removed from PBMC and CD34^+^ samples using Scrublet [81] algorithm by calculating doublet scores for each cell in each emulsion in the sample, clustering cells in high resolution using Spectral Clustering in scikit-learn package, evaluating mean doublet score and fraction of predicted doublets per cluster and removing clusters with doublet score and doublet fraction.

### UMAP construction and cell type annotation

The filtered matrices from PBMC and lung carcinoma (LC) samples were normalized to 10,000 total counts per cell (CP10k), log-transformed and scaled. After normalization, genes with at least 10 CP10k in at least 5 cells (10 cells for LC) were considered abundant and retained, mitochondrial and ribosomal genes were excluded and top 3000 genes for PBMC, top 2000 genes for LC samples, based on Fano factor [22] were used for PCA. Dataset integration was performed using scanpy.external.pp.harmony_integrate() function in scanpy package [82]. k-nearest neighbor graph was constructed using the adjusted principal components (number of nearest neighbors for PBMC samples k=20, for LC samples k=30) and was used to build UMAP representation. The resulting representation was used for exploration in interactive SPRING application [83]. For PBMCs, initial clustering using Spectral Clustering revealed a cluster (n=1,075) with lower complexity which was removed from further analysis. Final clustering for PBMCs was performed using the number of clusters k=10. Differential gene expression analysis (cluster vs rest of cells, Mann-Whitney U test with Benjamini-Hochberg correction) was performed and top 50 marker genes of each cluster (adjusted P value < 0.05) were used for manual cell type annotation (Supplementary Table S10). Markers to identify cell types included *MS4A1* and *CD79A* (B cells), *LEF1* and *CD8B* (CD8 T cells), *IL7R* and *LTB* (CD4 T cells), *GNLY*, *NKG7*, (NK cells), *LYZ*, *S100A8* (CD14^+^ Monocytes), *FCGR3A*, *LST1* (FCGR3A^+^ Monocytes), *CLEC10A*, *FCER1A* (cDC), *LILRA4*, *PLD4* (pDC), *PPBP*, *PLD4* (Megakaryocytes). To annotate cell types in the LC data, first clustering using PhenoGraph Leiden algorithm [84] with parameter resolution=2 revealed a cluster of lower complexity cells (n=1,670) with mitochondrial gene enrichment which were removed from further analysis. Following the removal of the cluster, a UMAP embedding was constructed again as described above. As initially, the graph was clustered using PhenoGraph Leiden implementation with resolution=2 and resulting clusters were assigned into myeloid, lymphoid or non-immune groups based on FACS CD45 status (positive/negative) and expression of canonical markers [7]. Finally, UMAPs were constructed as described above for each group separately (myeloid, lymphoid, epithelial/stromal) and clustered using PhenoGraph Leiden implementation with resolution=0.5 for myeloid, resolution=0.8 for lymphoid and resolution=0.6 for epithelial/stromal compartment. Within each group, differential gene expression analysis was performed as described above. In total, 38 cellular phenotypes were annotated based on marker genes (list is provided in Supplementary Tables S11-13) and the labels were then transferred to the original UMAP embedding containing all cells (n=32 937, Fig. 5e). For plotting purposes cells were grouped by broad phenotype labels (i.e. epithelial, endothelial, etc.) (Figure 5h, i).

### Cell trajectory reconstruction

To determine cell fate probabilities, we employed Palantir algorithm as described in Setty et al, 2019 [85]. To prepare the CD34^+^ data for Palantir, briefly, the filtered data was normalized, log-transformed and scaled as described above. Highly variable genes were selected using the scanpy highly_variable_genes() function with flavor argument set to ‘cell_ranger’. Cell cycle regression was performed using cell cycle genes defined in Tirosh et al, 2015 [86] as an input for scanpy score_genes_cell_cycle() function, first, to calculate cell cycle scores and then to regress out G2M and S scores using regress_out() function. Next, PCA was performed, data was integrated using scanpy.external.pp.harmony_integrate() function in scanpy package and k-neighbor graph (k=50) was constructed, as described above. The graph was used to build force-directed layout representation. Clustering and differential gene expression was performed as previously described above. Markers used for cell type annotation cell types were genes including *SPINK2*, *AVP* (HSC), *SPINK2*, *SMIM24* (MPP), *CD79B*, *VPREB1* (CLP), *ELANE*, *AZU1* (NMP), *KLF1*, *HBB* (Ery), *SCT*, *IRF8* (pDC), *PLEK*, *PPBP* (Mega), *LYZ*, *S100A9* (cDC), *CLC*, *PRG2* (Mast) (complete marker list is provided at Supplementary Table S14). Once data was prepared, Palantir package was used to construct diffusion maps using 10 components, and impute data using MAGIC. Early cell was determined in the HSC cluster employing the early_cell() function and was used as a starting point to run the Palantir algorithm with 1200 waypoints. Main differentiation trajectories of hematopoiesis were identified using the following markers: *CD34* for hematopoietic stem cells, *IRF8* for dendritic cells, *KLF1* for erythroid, *MYADM* for myeloid, *CD79A* for lymphoid and *ITGA2B* for megakaryocytes.

### Comparison of IVT– and TS-based inDrops-2

To calculate sequence alignment and quality control metrics, including gene-body coverages, mapping distributions across genomic features and read GC contents from the bam files, the ReSQC v5.0.1 [87] was employed with default parameters and GRCh38 genome (GENCODE v42) annotations. To compare gene length distributions between IVT– and TS-based protocols, gene lengths were retrieved from the .gtf annotation file (GENCODE v41, as described above) and were either binned into 10 or 3 categories, so that the number of genes is approximately equal in each of the category. The binning was performed using the ntile() function from dplyr v1.0.9. Filtered cell ⨉ gene expression data was size factor normalized and log_2_(x+1)-transformed using normalizeCounts() with default parameters from the scuttle v1.6.3. Fractions of each gene length category per cell were calculated by dividing the sum of counts per category by the total counts. Differential gene expression analysis between IVT– and TS-based scRNA-Seq data was performed using the MAST package [88]. More explicitly, hurdle models were fitted on filtered, size factor normalized and log_2_(x+1)-transformed expression data using protocol type and number of detected genes (centered) as covariates to adjust for the cellular detection rate. Gene set enrichment analysis (GSEA) was performed with a sorted gene list (by descending log2 fold change values) and gene ontology (GO) [89] terms using the gseGO() function from ClusterProfiler package [90]. The resulting P values were adjusted using Benjamini-Hochberg FDR correction.

